# Single-cell analyses reveal a continuum of cell state and composition changes in the malignant transformation of polyps to colorectal cancer

**DOI:** 10.1101/2021.03.24.436532

**Authors:** Winston R. Becker, Stephanie A. Nevins, Derek C. Chen, Roxanne Chiu, Aaron Horning, Rozelle Laquindanum, Meredith Mills, Hassan Chaib, Uri Ladabaum, Teri Longacre, Jeanne Shen, Edward D. Esplin, Anshul Kundaje, James M. Ford, Christina Curtis, Michael P. Snyder, William J. Greenleaf

**Author notes:** Correspondence (M.P.S.), (W.J.G.). These authors contributed equally (Winston R. Becker, Stephanie A. Nevins).

## Abstract

To chart cell composition and cell state changes that occur during the transformation of healthy colon to precancerous adenomas to colorectal cancer (CRC), we generated 451,886 single-cell chromatin accessibility profiles and 208,557 single-cell transcriptomes from 48 polyps, 27 normal tissues, and 6 CRCs collected from patients with and without germline APC mutations. A large fraction of polyp and CRC cells exhibit a stem-like phenotype, and we define a continuum of epigenetic and transcriptional changes occurring in these stem-like cells as they progress from normal to CRC. Advanced polyps contain increasing numbers of stem-like cells, regulatory T-cells, and a subtype of FOX-regulated pre-cancer associated fibroblasts. In the cancerous state, we observe T-cell exhaustion, RUNX1-regulated cancer associated fibroblasts, and increasing accessibility associated with HNF4A motifs in epithelia. Methylation changes in sporadic CRC are strongly anti-correlated with accessibility changes along this continuum, further identifying regulatory markers for molecular staging of polyps.

## INTRODUCTION

The identification of key genes and pathways that drive the formation of invasive cancers has been the central focus of a number of large-scale genomics efforts (Consortium and The ICGC/TCGA Pan-Cancer Analysis of Whole Genomes Consortium, 2020; International Cancer Genome Consortium et al., 2010; Weinstein et al., 2013). These efforts have successfully cataloged the diversity and commonality of many of the genetic and transcriptional changes that accompany malignancy in diverse cancer types. However, most of this work has focused on bulk profiling of advanced stage tumors, and has largely ignored premalignant lesions. As a result, a detailed understanding of the progression of phenotypic changes that occur during the transition from normal to precancerous to cancerous state, as well as the molecular drivers of this transformation, remain under-explored.

Colorectal cancer (CRC) is an ideal system to study the continuum of phenotypic states along malignant transformation as it follows a stereotyped progression from normal to atypical to carcinoma. The steps along this pathway include the formation of precancerous polyps that are readily identifiable and thus suitable for isolation and study (Aoki and Taketo, 2007; Fodde et al., 2001). These polyps can subsequently give rise to CRCs, and a number of the changes associated with these transitions are nearly universal to all CRC malignancies, as typified by the adenoma-to-carcinoma sequence (Fearon and Vogelstein, 1990; Leslie et al., 2002; Vogelstein et al., 1988). For example, an estimated 80-90% of colorectal tumors are initiated by loss of APC, a member of the WNT signaling pathway and β-catenin complex (Mori et al., 1992). Loss of APC results in β-catenin stabilization and increased WNT signaling (Logan and Nusse, 2004), leading to intestinal hyperplasia (Schatoff et al., 2017). Subsequent mutations in other cancer driver genes such as *KRAS, TP53,* and *SMAD4* result in the transformation to carcinoma.

Because *APC* mutations are almost universally the initiating event for polyps and CRCs, Familial Adenomatous Polyposis (FAP) patients, who have germline mutations in *APC*, are an ideal population in which to study the natural progression of polyposis. These patients typically develop hundreds of polyps in adolescence or early adulthood (Galiatsatos and Foulkes, 2006; Groden et al., 1991), and therefore an individual patient can provide a source of numerous polyps of varied molecular ages and stages of progression, all arising in the same germline genetic background.

To chart the regulatory and transcriptomic changes that occur on the phenotypic continuum from healthy colon to invasive carcinoma, as part of the Human Tumor Atlas Network (HTAN) (Rozenblatt-Rosen et al., 2020), we profiled single-cell transcriptomes (snRNA-seq) and epigenomes (scATAC-seq) of healthy colon, polyps, and CRCs. Many of the polyps were obtained from FAP patients who underwent surgical colectomies, a procedure involving partial or complete removal of the colon. Targeting this colectomy tissue for analysis allowed us to 1) identify polyps with diverse sizes and locations of origin, and 2) collect neighboring unaffected colon tissue. From the single-cell datasets generated from these samples, we first catalogue immune, stromal, and epithelial cell types in normal colon, polyps, and CRC. We find large shifts in fibroblast subpopulations that occur along the transition from normal colon to CRC. We also identify a subpopulation of exhausted T-cells, which do not appear in normal tissues or polyps, but only in CRC tissue. We observe a much larger fraction of cells in polyps exhibit a stem-like state (both transcriptionally and epigenetically) within the polyp tissues. We find that polyps populate an epigenetic and transcriptional continuum from normal colon to CRC characterized by sequential opening and closing of chromatin and upregulation and downregulation of genes associated with the cancer state. Furthermore, the fraction of cells within a polyp exhibiting a stem-like state is an accurate proxy for how close a polyp is to the molecular phenotypes associated with carcinoma. We identify regulatory elements and transcription factors associated with the different stages of transformation from normal colon to carcinoma. Early changes in regulatory elements are characterized by increased accessibility of peaks containing TCF and LEF motifs and loss of accessibility of peaks containing KLF motifs. In the final stage of this pathway, malignant transformation, we observe an increase in accessibility of peaks containing HNF4A motifs. Finally, we show that accessibility changes in polyps are strongly anti-correlated with methylation changes in sporadic CRC, and demonstrate that a subset of loci that exhibit methylation changes associated with cancer change their accessibility state very early in the malignant continuum, suggesting potential strategies for the detection of premalignant polyps.

## RESULTS

### MAPPING MOLECULAR CHANGES ACROSS THE CONTINUUM OF MALIGNANT TRANSFORMATION

We generated single-cell data for a total of 81 samples collected from 15 donors (8 FAP and 7 non-FAP) (Figure 1A, Supplemental Tables S1 and S2). These data include deep profiling of 4 FAP patients from whom we assayed 8–11 polyps, 0–1 carcinomas, and 4–5 matched normal (unaffected) tissues. From non-FAP donors, we collected data on normal colon (9 samples from 2 donors), polyps (1 sample from 1 donor), and colorectal cancer (CRC) tissues (4 samples from 4 patients). For each tissue, we performed matched scATAC-seq and snRNA-seq (10x Genomics). Overall, we obtained high quality single-cell chromatin accessibility profiles for 451,886 cells from 80 samples, with a mean TSS enrichment of approximately 8 for most samples (Figure S1A). After removing low quality snRNA-seq cells and samples, we obtained single-cell transcriptomes for 208,557 cells from 72 samples (Figure S1B).

**Figure 1:**
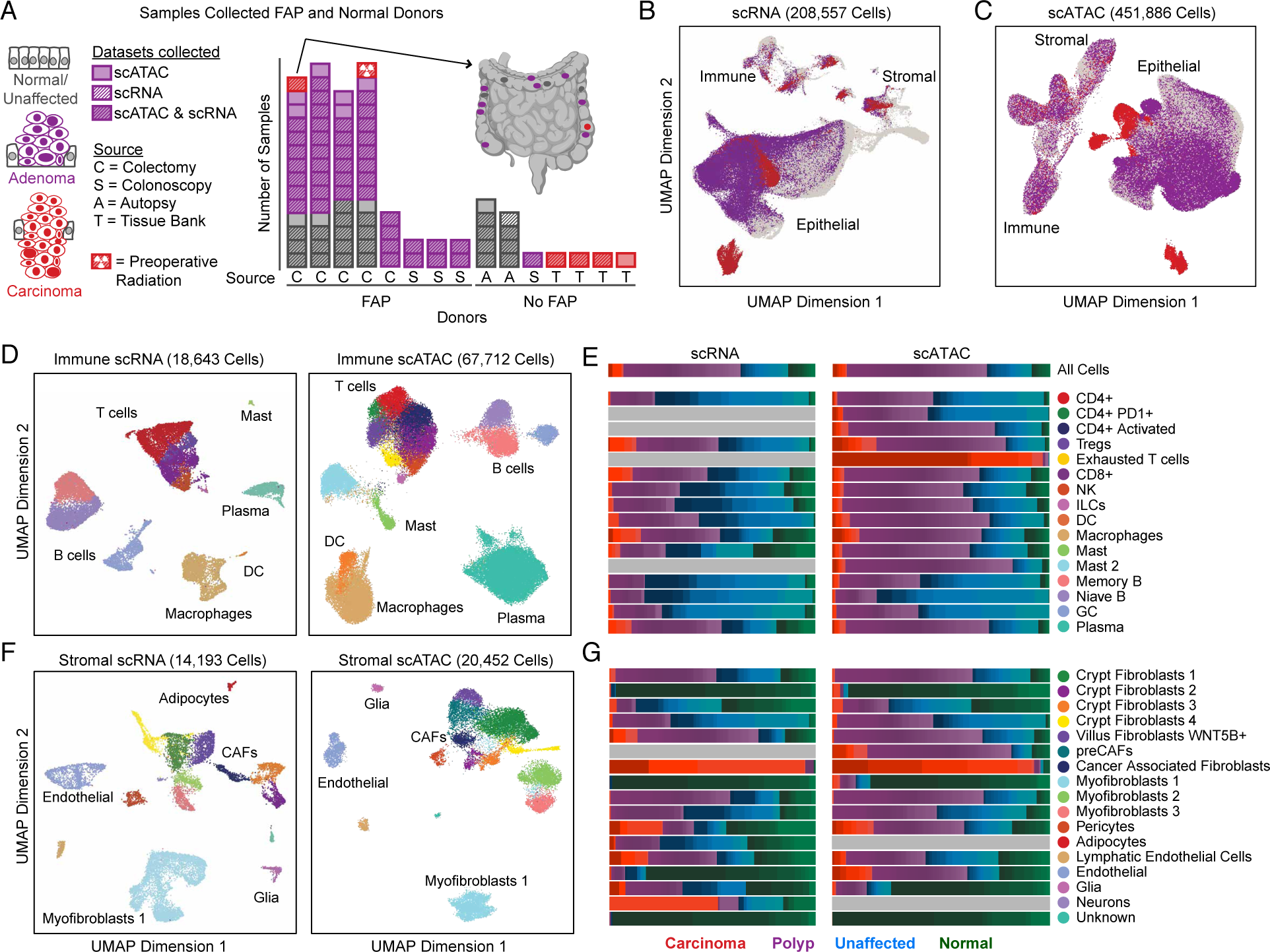
Multi-omic single-cell atlas of cancer development. (A) Summary of the samples in this study. The bar chart shows the number of normal/unaffected colon tissues (gray), adenomas (purple), and CRCs (red) assayed for each patient. Locations of samples assayed from one patient are indicated on the colon on the upper right. (B, C) UMAP representations of all snRNA-seq (B) and scATAC-seq (C) cells colored by whether the cells were isolated from normal/unaffected colon tissues, adenomas, or CRCs. (D,F) UMAP representations and annotations of immune (D) and stromal (F) cells. (E,G) Fraction of each immune (E) and stromal (G) cell type isolated from normal (green), unaffected (blue), polyp (purple), and CRC (red) samples.

When all snRNA-seq cells (Figure 1B) and scATAC-seq cells (Figure 1C) are projected into low dimensional subspaces, stromal and immune cells generally cluster by cell type whereas epithelial cells largely separate into distinct clusters comprising cells derived from polyps, unaffected tissues, or CRCs. As a result, we were able to annotate immune and stromal cells by sub-clustering cells from all samples and analyzed the epithelial subset separately.

### ANNOTATION OF IMMUNE AND STROMAL CELL TYPES REVEALS ENRICHMENT OF T-CELLS AND MYELOID CELLS IN POLYPS AND CRC

The immune compartment derived from these tissues consisted of multiple clusters of B-cells and T-cells, as well as monocytes, macrophages, dendritic cells (DCs), and mast cells (Figure 1D). To annotate the snRNA cells, we examined gene expression of known marker genes (Figure S1C). Similarly, to annotate the scATAC cells, we examined chromatin activity scores— a measure of accessibility within and around a given gene body—associated with these marker genes (Figure S1D). As expected, clusters of B-cells exhibited both high *PAX5* chromatin activity scores and gene expression, clusters of plasma cells exhibited high *IGLL5* activity scores and expression, and clusters of regulatory T-cells (Tregs) exhibited high *FOXP3* activity scores and expression (Figure S1C,S1D).

We validated our manual snRNA-seq cluster annotations with automated cell annotation software (singleR; Figure S1E) (Aran et al., 2019). Although singleR annotations are limited by the reference dataset, the most common cell types in each cluster generally agreed with our manual annotation (Figure S1E). The cell types identified were represented by nearly all samples, although some cell types were notably enriched or depleted in specific disease states (Figure 1E). For example, a larger fraction of the myeloid, Tregs, and exhausted T-cells came from polyps or CRCs, while a larger fraction of Memory B, Naive B, and germinal center (GC) B cells came from normal and unaffected tissues. This enrichment of myeloid and specific types of T-cells and dis-enrichment of B-cells has been recently reported in CRC (de Vries et al., 2020), and thus we observe that these shifts in the tumor immune composition also occur in precancerous polyps.

Within the stromal compartment, we identified glial cells, adipose cells, and multiple types of endothelial cells and fibroblasts (Figure 1F). Fibroblasts within the normal colon are known to participate in variable degrees of WNT and BMP signaling to support the functions of the colonic mucosa (Powell et al., 2011; Smillie et al., 2019). Within our single-cell datasets, we identify 10 clusters of fibroblasts. Subtypes of fibroblasts include those present at the crypts, characterized by expression of *WNT2B* and *RSPO3*, fibroblasts thought to be present at villi, characterized by expression of *WNT5B*, and myofibroblasts, which express high levels of *ACTA2* and *TAGLN* (Figure S2A) (Powell et al., 2011). These fibroblast subtypes are observed in normal/unaffected tissues as well as polyps (Figure 1G). As previously shown, villus fibroblasts play important roles in BMP signaling, and BMP genes are highly expressed in villus fibroblasts in our dataset (Figure S2A). Recent reports also found that crypt fibroblasts secrete semaphorins to support epithelial growth, and we observe one cluster of fibroblasts with high expression of semaphorins (Figure S2A) (Karpus et al., 2019). This same cluster of fibroblasts exhibited the highest expression of *RSPO3*, a factor that supports the intestinal stem cell niche. We also observe a cluster of cancer associated fibroblasts (CAFs) that consists almost exclusively of cells from CRCs, and a scATAC cluster of fibroblasts with accessibility around many of the same genes as the CAFs, which we term pre-cancer associated fibroblasts (preCAFs), and is enriched for cells from both CRCs and polyps (Figure 1G). These observations suggest that phenotypically distinct fibroblasts exist in polyps as well as tumors, and thus may play a role in tumorigenesis in precancerous lesions.

After annotation of the immune and stromal compartments, we integrated our scATAC and snRNA datasets to enable additional analyses of regulatory elements and transcription factors potentially driving the expression changes. We linked our scATAC and snRNA data by aligning the datasets with CCA and assigning RNA-seq profiles to each scATAC-seq cell (Granja et al., 2019), then labeled the scATAC cells with the nearest snRNA cells. This labelling closely agreed with our manual annotations of the immune (Figure S1I) and stromal (Figure S2B) subcompartments. To nominate potential peak-to-gene links, we identified peaks that were highly correlated to gene expression in our datasets. Within the stromal compartment, this analysis enabled identification of 52,443 peak-to-gene links (correlation≥0.45; Figure S2D).

### SINGLE-CELL CHROMATIN ACCESSIBILITY PROFILING REVEALS POPULATIONS OF CANCER SPECIFIC FIBROBLASTS AND FIBROBLASTS BECOMING CANCER LIKE

CAFs promote cancer development and progression through diverse mechanisms including matrix remodeling, signaling interactions with cancer cells, and perturbation of immune surveillance (Karagiannis et al., 2012; Koliaraki et al., 2017; Tommelein et al., 2015). In our data, we observe a CAF cluster composed entirely of cells derived from adenocarcinoma, with effectively no contributions from polyp, unaffected, or normal tissue. Cells in this cluster have high expression of known CAF marker genes *FAP* and *TWIST1* (Figure S2A) (Lee et al., 2015; Puré and Blomberg, 2018). We identified marker genes most significantly associated (Wilcoxon test) with the CAF cluster in the snRNA-seq data (Figure 2C). Among the most specific markers for CAFs compared to other fibroblast subtypes were 1) FAP, a serine protease that is upregulated in fibroblasts in a number of cancers (Puré and Blomberg, 2018), 2) VCAN, which encodes the proteoglycan versican that is implicated in growth and metastases of several cancer types, and is proposed to promote angiogenesis (Asano et al., 2017), and 3) COL1A2, a subtype of type I collagen highly expressed in many cancers and implicated in extracellular matrix remodeling in malignancy (Fang et al., 2019). Specific expression of these genes by CAFs in CRC suggests that fibroblasts participate in unique extracellular matrix remodeling in cancerous tissues that does not occur in normal colon or precancerous polyps.

**Figure 2:**
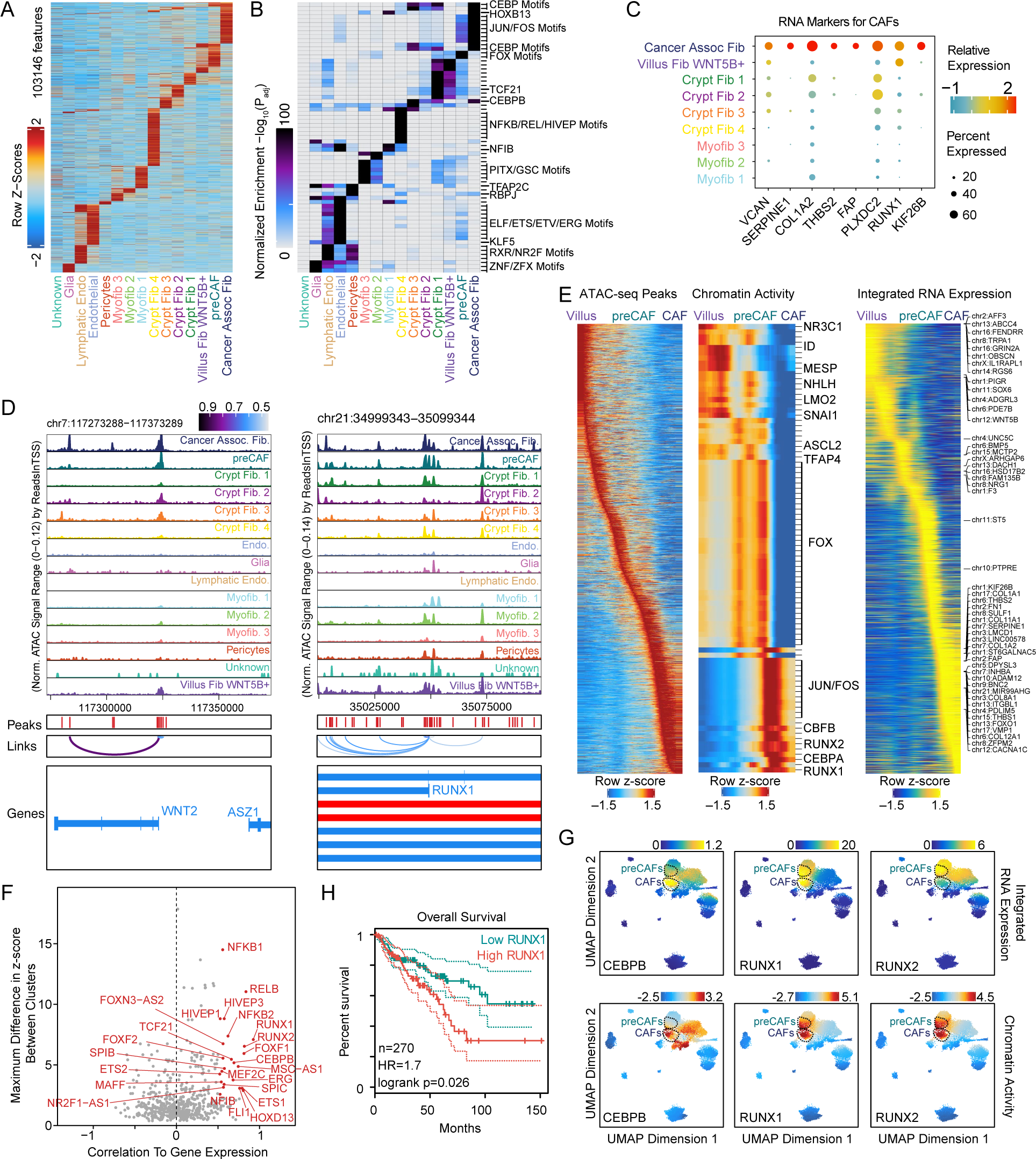
Epigenetic regulators of pre-cancer-associated and cancer-associated fibroblasts. (A) Marker peaks (wilcoxon FDR ≤ 0.1 & Log_2_FC ≥0.5) for each stromal cell type. (B) Hypergeometric enrichment of transcription factor motifs in stromal cell marker peaks. (C) Dotplot representation of the most significant (wilcoxon test) marker genes for CAFs. (D) Genomic tracks for accessibility around *WNT2* and *RUNX1* for different stromal cell types. Peaks called in the scATAC data and significant peaks-to-gene links are indicated below the tracks. (E) Changes in most variable peaks, TF motif activity scores, and gene expression along the trajectory from villus fibroblasts to preCAFs to CAFs. (F) Plot of maximum difference between chromVAR deviation z-score, depicting TF motif activity, against correlation of chromVAR deviation and corresponding TF expression. TF’s with maximum differences in chromVAR deviation z-score in the top quartile of all TF and a correlation of greater than 0.5 are indicated in red. (G) RNA expression (top) and chromVAR deviation z-scores (bottom) for selected transcription factors. The RNA expression plotted is the expression in the nearest RNA cell following integration of the snRNA and scATAC data. (H) Overall survival of colon adenocarcinoma TCGA patients with expression of RUNX1 in the top and bottom half of the group (log-rank test).

While CAFs are known to promote colon cancer progression, and we observe a subset of CAFs unique to CRCs, we next explored the role of fibroblasts in precancerous lesions. The fibroblast cluster most enriched for cells from polyps was the preCAF cluster identified in the scATAC data. When we examined accessibility around marker genes for CAFs we found that many of them were also more accessible in the preCAF cluster than other fibroblast subtypes. For example, CAFs secrete Wnt2 to promote cell proliferation and cancer progression (Aizawa et al., 2019) as well as angiogenesis in CRC (Unterleuthner et al., 2020). CAFs and preCAFs exhibit the greatest accessibility at the *WNT2* TSS (Figure 2D), suggesting that chromatin changes promote the expression of *WNT2* in CAFs and preCAFs. When we examine which peaks are most correlated to the expression of *WNT2*, we identify a regulatory element ∼50KB away from the TSS that is primarily only accessible in CAFs, and is potentially driving higher expression of *WNT2* in these cells. However, the observation that, when compared to other populations of fibroblasts, preCAFs had relatively higher accessibility at many known CAF marker genes led us to hypothesize that these preCAFs may be on the path towards becoming CAFs.

We next asked which transcription factors are associated with differential accessibility between fibroblast subtypes. We identified marker peaks (Wilcoxon FDR≤0.1; Log_2_FC≥0.5) for each cell type in the stromal compartment relative to a background of all other cell types (Figure 2A) and then computed the hypergeometric enrichment of motifs within marker peaks for each cell type (Figure 2B; Methods). Most stromal clusters have specific transcription factor enrichment in their marker peaks, suggesting differential and relatively distinct regulation occurs in each of these cell types. Notably, marker peaks for CAFs were enriched for JUN/FOS and CEBP motifs. In preCAF marker peaks we also observed enrichment of JUN/FOS motifs, but to a lesser extent, as well as enrichment of both FOX motifs and motifs enriched in Villus and Crypt fibroblasts. The overlap between motifs enriched in marker peaks for the preCAF and CAF clusters further supports the hypothesis that preCAFs represent an intermediate cell state between normal colon fibroblasts and CAFs.

To further map out the changes in these populations of fibroblasts, we constructed a trajectory between villus fibroblasts, preCAFs, and CAFs and examined changes in peaks and chromVAR activity scores along this putative trajectory. When we plot the most variable peaks, we observe a relatively monotonic opening and closing of peaks along this trajectory (Figure 2E). Chromatin accessibility activity levels of DNA motifs (quantified using chromVAR deviation z-scores) along this trajectory show increased activity of FOX family transcription factors in the intermediate states, that is followed by increased activity of JUN, FOS, CEBP, and RUNX1 motifs in CAFs (Figure 2E). The smooth accessibility and expression changes along this putative trajectory are consistent with the hypothesis that normal colon fibroblasts, such as villus fibroblasts, develop into preCAFs and eventually CAFs.

### RUNX1 IS A KEY TRANSCRIPTION FACTOR ASSOCIATED WITH WIDESPREAD CHROMATIN ACCESSIBILITY AND GENE ACTIVITY IN CAFs

The above analysis highlighted TF motifs that are associated with chromatin accessibility in different populations of stromal cells. However, many TFs share similar motifs, so identification of the precise factors that are functional in these cell types is aided by also examining TF expression. To nominate TFs driving changes in chromatin accessibility in different stromal cell types, we identified TFs with the highest correlation between their gene expression and the chromatin accessibility activity level of the DNA motif it was expected to bind (Figure 2F, x-axis). Amongst the most correlated TFs were RUNX1, RUNX2, and CEBPB. We next plotted the expression and motif activities of these TFs on the UMAP representation of the stromal cells (Figure 2G), and noted that chromatin activity levels for RUNX1 and RUNX2, which have similar motifs, are highest in CAFs and preCAFs. However, *RUNX1* is primarily expressed in CAFs and preCAFs, while *RUNX2* has much lower expression in CAFs. This suggests that RUNX1 is a stronger driver of accessibility at RUNX motifs than is RUNX2 in CAFs.

We next inspected chromatin accessibility near *RUNX1* to identify regulatory elements that may be driving increased RUNX1 activity and, consistent with the expression of these genes, observed the greatest accessibility around the TSS in CAFs and preCAFs (Figure 2D). We identify a number of regulatory elements whose accessibility is correlated with the expression of *RUNX1* in our linked snRNA dataset (Figure 2D). Given that RUNX1 is responsible for some CAF specific gene expression, we examined if expression of RUNX1 was related to overall survival in patients with CRC using previously published TCGA data and found that patients with higher expression of RUNX1 had significantly worse overall survival, possibly because this correlates with increased presence of CAFs driving tumor progression (Figure 1H).

### EXHAUSTED T-CELLS ARE PRESENT IN CRC, BUT NOT IN PRECANCEROUS ADENOMAS

A primary immune function is to recognize and destroy tumor cells to prevent cancer formation. However, multiple processes of immune evasion, including immune suppression by Tregs cells, defective antigen presentation, production of immunosuppressive cytokines, and immune cell dysfunction, allow tumors to overcome immune surveillance (Vinay et al., 2015). We aimed to determine if there is evidence of immune evasion in precancerous lesions, and if so, when in the malignant transformation trajectory it occurs.

T-cell exhaustion, a dysfunctional state that occurs in response to chronic antigen stimulation, is characterized by reduced cytokine production and increased expression of inhibitory receptors and is thought to be a primary mechanism of immune evasion by cancers (Blank et al., 2019; Jiang et al., 2015). We aimed to determine if T-cell exhaustion could be detected in early stages of the transition to carcinoma or only later after malignant transformation. We sub-clustered the T-cells in our data set, projected these cells into a UMAP, and colored the cells by disease-state of the tissue of origin or their cell type annotations (Figure 3A,B). We next identified exhausted T-cells in four different ways. First, we examined gene activity scores for exhausted T-cells markers including *BATF*, *CTLA4*, *PDCD1*, and *TOX*, and found that they were high in one cluster (Figure 3C). Second, we used a previously published snRNA-seq dataset that contained exhausted T-cells from basal cell carcinoma (Yost et al., 2019) to identify exhausted T-cells within the sub-clustering of T-cells by aligning the datasets with CCA and labeling our cells by the closest RNA-seq profiles as described above (Yost et al., 2019) (Figure 3E). Third, we identified exhausted T-cell specific regulation by identifying differentially accessible peaks (Wilcoxon test) relative to CD8+ T-cells (Figure 3F). Peaks more accessible in exhausted T-cells were enriched for NR4A, RUNX, CEFB, JUN, FOS, and BATF family motifs, many of which are known drivers of T-cell exhaustion (Satpathy et al., 2019). Fourth, we computed chromVAR deviations, which also show that BATF and NR4A2 motifs tend to be more accessible in exhausted T-cells (Figure 3D). Together, these four lines of evidence identified the same population of exhausted T-cells, which are observed in CRC samples only, demonstrating that this specific immunological dysfunction seems to be unique to invasive cancer samples.

**Figure 3:**
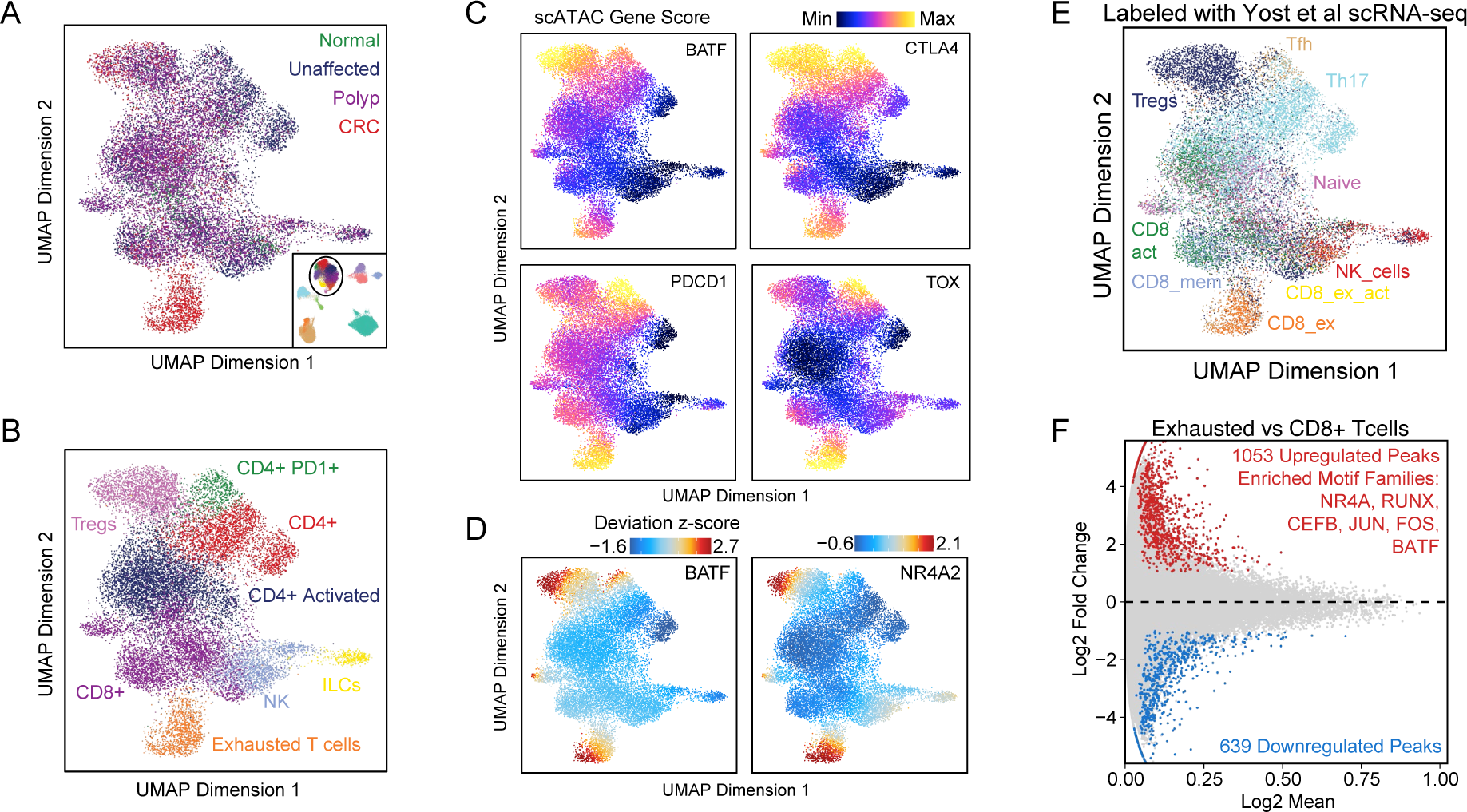
Exhausted T-cells are specific to CRC. (A, B) UMAP projection of all T-cells identified in the scATAC data. Points on the UMAP represent single-cells and are colored by tissue of origin (A) and cell-type annotations (B). (C) UMAP projection of scATAC T-cells with cells colored by gene activity scores depicting chromatin accessibility surrounding *BATF*, *CTLA4*, *PDCD1*, and *TOX*. (D) ChromVAR deviation z-scores depicting TF motif activity of BATF and NR4A2 plotted on scATAC T-cell UMAP. (E) UMAP projection of scATAC T-cells colored by labeling of scATAC-seq T-cells with nearest snRNA-seq T-cells in BCC after integrating the datasets with CCA. (F) MA plot showing differential peaks (wilcoxon test) between exhausted T-cells and CD8+ T-cells. Motifs with hypergeometric enrichment in peaks more accessible in exhausted T-cells are listed in the plot.

### POLYPS ARE ENRICHED FOR EPITHELIAL CELLS OCCUPYING A STEM-LIKE STATE

We next examined in detail the epithelial cells in which unaffected, polyp, or CRC initially clustered largely by disease state (Figures 1B and 1C). To better analyze epithelial cells in different disease subtypes, we first constructed an RNA and ATAC-seq reference map of normal epithelial colon cells collected from genetically normal patients (Figure 4A). We annotated cell types in this normal tissue using gene expression and gene activity scores of known marker genes (Figure S3A). A stem cell population with high expression and accessibility of *LGR5* and *SMOC2* was immediately evident, as were goblet cells with high expression of *MUC2*, and Best4+ enterocytes exhibiting high expression of *BEST4*. Following manual annotation of these clusters, the snRNA and scATAC datasets were aligned with CCA (Granja et al., 2019; Stuart et al., 2019), and the scATAC cells were labeled based on the nearest snRNA cells, which produced excellent agreement (65% accuracy overall) with the manual annotations, with mislabeled cells typically being labeled as the nearest cell type in the differentiation trajectory (Figures S3C and S3D).

**Figure 4:**
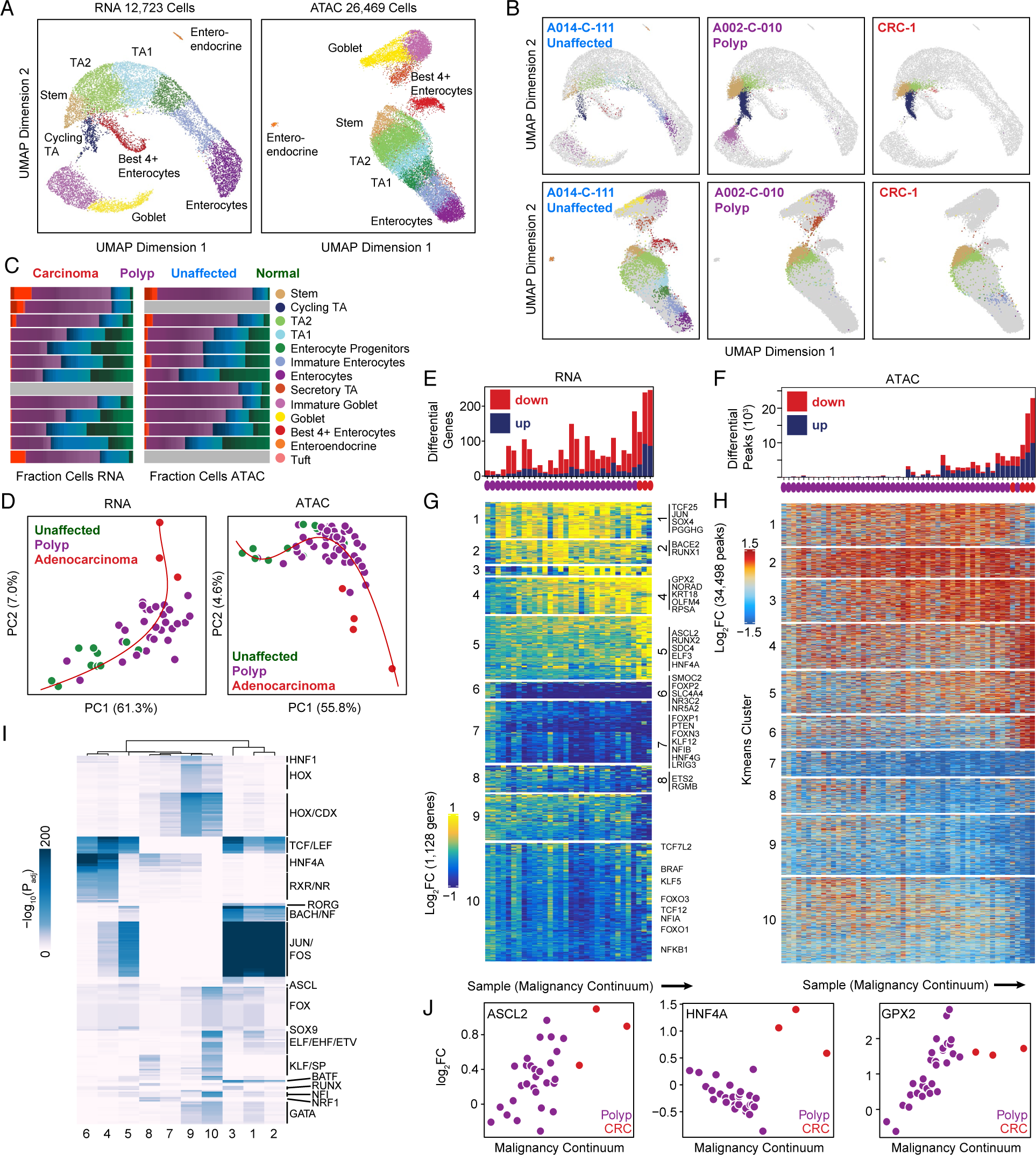
The regulatory trajectory of malignant transformation. (A) UMAP projection of snRNA (left) and scATAC (right) epithelial cells isolated from normal colon with cells colored by cell type. (B) Projection of epithelial snRNA (top) and scATAC (bottom) cells from unaffected (left), polyp (center), and CRC (right) samples into the manifold of normal colon epithelial cells. Projected cells are colored by nearest normal cells in the projection and normal epithelial cells are colored gray. (C) Fraction of each epithelial cell type isolated from normal (green), unaffected (blue), polyp (purple), and CRC (red) samples. Cell types are defined based on the identity of the nearest cell types when projecting epithelial cells into normal colon subspace. (D) Malignancy continuum for snRNA (left) and scATAC (right). Principle components were computed on the log_2_ fold changes between stem-like cells from each sample and normal colon stem cells for the set of peaks and genes that were significantly differential (wilcoxon test) in at least 2 samples. A spline was fit to the first two principal components (red) and samples were ordered based on their position along the spline. (E, F) Number of significantly differential (wilcoxon test) genes (E) and peaks (F) for each sample relative to all unaffected samples. (G, H) Heatmap of all genes (G) and peaks (H) that were significantly differentially expressed (wilcoxon test, p_adj_≤0.05 & |log_2_FC|≥0.75) or accessible (wilcoxon test, p_adj_≤0.05 & |log_2_FC|≥1.5) in ≥2 samples. Samples are ordered along the x-axis by the malignancy continuum defined in D. Genes and peaks are k-means clustered into 10 groups. (I) Hypergeometric enrichment of TF motifs in k-means clusters of peaks defined in H. (J) Log_2_FC in expression of *ASCL2*, *HNF4A*, and *GPX2* in stem-like cells from each sample relative to stem-like cells in unaffected samples plotted against the malignancy continuum defined in D. Samples are colored based on if they are derived from polyps or CRCs.

To determine the gene expression and epigenetic state of FAP unaffected, polyp, and CRC samples in the context of normal tissue, we projected these cells into this normal subspace, then classified each cell based on the nearest normal epithelial cell (Figure 4B) (Granja et al., 2019). This projection immediately revealed that epithelial cells from polyps and CRCs tend to project closer to stem cells and other immature cells along the normal differentiation trajectory, whereas cells from unaffected tissues projected relatively evenly throughout the epithelial compartment (Figure 4B). We classified all epithelial cells based on the nearest normal cells in the projection and find that cells originating from polyps and CRC samples are enriched for stem-like epithelial cells and dis-enriched for mature enterocytes, suggesting that epithelial cells increasingly demonstrate a stem-like phenotype during the transformation from normal to polyp (Figure 4B, 4C).

To further characterize these stem-like cells observed in polyps and CRCs, we examined expression of intestinal stem cell and colon cancer stem cell marker genes in these stem-like populations. A number of markers for intestinal stem cells have been identified, including LGR5, SMOC2, RGMB, SOX9, and LRIG1 (Heijden et al., 2019). Colon cancer stem cells have also been reported to express LGR5, and have been proposed to be CD133+, EPCAM^high^, CD44+, CD166+, ALDH+, and EphB2^high^ (Heijden et al., 2019). Additionally, ASCL2 has been shown to be a master regulator for intestinal stem cells (van der Flier et al., 2009). Among these markers, we find that *LRIG1* and *RGMB* are expressed primarily in stem cells from normal and unaffected tissues, *LGR5*, *EPHB2*, *ALCAM*/*CD166* are highly expressed in stem-like cells from normal, unaffected, polyp, and CRC samples, and *CD44*, *EPCAM*, *ASCL2*, and *SOX9* are highest expressed in stem-like cells from polyps and CRCs (Figure S4). *LGR5* expression is specific to stem-like cells in all samples, and has low or no expression in other cell types, further supporting that projecting diseased cells into the normal manifold effectively identified stem cell populations.

### STEM-LIKE CELLS ISOLATED FROM NORMAL COLON, POLYPS, AND CRC EXHIBIT A CONTINUOUS TRAJECTORY OF DIFFERENTIAL EXPRESSION AND REGULATION AND FORM A POTENTIAL MALIGNANCY CONTINUUM

The above analysis suggests that polyps and CRCs have a large population of cells that are most similar to normal stem cells. We next wanted to determine if, in addition to these changes in cell composition, the cell state of stem-like cells changes in precancerous and cancerous lesions. To identify these differences in cell state, we compared the gene expression and chromatin accessibility of polyp and CRC stem-like cells to normal stem cells. By making this comparison to the nearest normal cell type we aimed to identify the aberrant gene expression and regulatory programs occurring in these stem-like cells after “subtracting” the normal stem-like programs observed in wild type stem cells.

After computing differential peaks (Wilcoxon FDR≤0.05; |Log_2_FC|≥1.5) between stem-like cells from each sample and cells from the nearest normal cell type, we projected our samples into the first two principal components of signals from these differential peaks and ordered samples based on their position along a spline fit in this space (Figure 4D). We generated a similar trajectory for the RNA data using differential genes rather than differential peaks (see Methods). The ordering of samples along the continuums defined from the RNA and ATAC datasets exhibited strong agreement overall (Figure S3H). This analysis suggests that the differences in gene expression and chromatin accessibility between stem cells and these stem-like cells follow a stereotyped progression from early to late polyp to invasive CRC.

The order of the stem cells may serve as a proxy for the progression of a polyp toward malignant transformation. Nearly all of the polyps in our study were graded as containing low-grade dysplasia; however, our molecular findings suggest these polyps represent a large diversity of phenotypes on a relatively continuous spectrum from normal epithelia to invasive cancer. We found that samples with no dysplasia on microscopic pathology are the earliest samples on the malignancy continuum. Notably, two samples that we classified as polyps during sample collection were found to have no dysplasia on pathology review, and these samples were the earliest polyps in the malignancy pseudotime. Further, two unaffected samples were found to have low grade dysplasia, and these unaffected samples were the unaffected samples the furthest along the malignancy continuum. Beyond this binary separation of samples with dysplasia or no dysplasia, the degree of transformation in each polyp was not highly correlated with the fraction of cells classified as dysplastic as determined from microscopic pathology (Figure S3G), although this dysplastic fraction measure is not used as a clinically relevant metric for assessing polyp severity. This lack of correlation with microscopic pathology suggests that molecular phenotyping provides a higher resolution as to the degree of malignant transformation than does traditional H&E staining and microscopy-based pathology review. We also note that the pathology measurements and single-cell assays are performed on different sections of the same polyp, so we cannot rule out that some differences may be attributed to heterogeneity between sections of the same polyp. While we cannot currently assess how clinically meaningful these differences may be, the presence of high-grade vs. low-grade dysplasia is currently used as a criteria for timing of repeat colonoscopy, and we therefore speculate that our malignancy metric may also be useful for risk stratification and colonoscopy timing.

After computing the trajectory, and given that we observed relatively few differences between the FAP unaffected colon and normal colon, we repeated the differential analysis using all unaffected samples rather than normal samples to increase the total number of patients and cells in the background group. We observe that the absolute number of significantly differential peaks and genes gradually increased along the malignancy continuum—with adenocarcinoma samples exhibiting the largest number of differential peaks and genes (Figure 4E and 4F).

### GENES INVOLVED IN OXIDATIVE STRESS RESPONSE, WNT SIGNALING, AND CELL MOTILITY ARE UPREGULATED ALONG THE MALIGNANT CONTINUUM

We next examined gene expression changes along this malignancy continuum. We selected genes differentially expressed in at least two samples and clustered these differentially expressed genes into 10 k-means clusters. These clusters correspond to groups of genes that become differentially expressed at distinct stages of malignant transformation. For example, clusters 1–4 consist of genes that become upregulated in stem-like cells in early stage polyps when compared to unaffected stem cells. Members of cluster 4 include *OLFM4*, a marker of intestinal stem cells (Flier et al., 2009), indicating that *OLMF4* is not just a stem-cell marker, but becomes even higher expressed in stem-like cells from polyps as they approach malignancy. Members of cluster 4 also include GPX2, a glutathione peroxidase known to be upregulated in CRC that functions to relieve oxidative stress by reducing hydrogen peroxide, facilitating both tumorigenesis and metastasis (Emmink et al., 2014). *GPX2* is upregulated even in early stage polyps, with its expression increasing across the malignancy continuum until plateauing in the CRC samples (Figure 4J). Further, the upregulation is not donor dependent, and we observe the same trend across all donors in our study (Figure S3D). We therefore speculate that the need to combat increased levels of oxidative stress occurs early in tumorigenesis.

Multiple gene clusters gradually reduce expression in the transition from normal colon to cancer (clusters 6–9). Members of these clusters include *NR3C2*, which codes for the mineralocorticoid receptor (Figure S3D). Low expression of *NR3C2* has been observed in multiple cancers and is associated with poor prognosis. Knockdown of *NR3C2* in hepatocellular carcinoma cells leads to an increase in β-catenin expression (Yang et al., 2019). Therefore, decreased expression of *NR3C2* in CRC may be another means to increase expression of WNT target genes and drive proliferation in polyps and CRC. LRIG3, a transmembrane protein recently reported to repress cell motility and metastasis in CRC (Zeng et al., 2020) also gradually decreases in expression across the malignancy continuum (Figure S3D).

While we observe many genes that are gradually upregulated or downregulated along the malignancy continuum, we also wanted to highlight genes specific to malignant transformation as they may represent the transcriptomic alteration necessary for invasion (Figure S4H). Many of the genes specific to malignant transformation are previously described markers for CRC that we find to be specific to CRC and not highly expressed in adenomas. For example, BMP7, a secreted signaling factor, is specifically upregulated in CRC and has been shown to correlate with metastasis and poor prognosis in CRC (Motoyama et al., 2008). Another gene found to be specifically expressed in CRC was DPEP1, a zinc-dependent metalloprotease involved in glutathione and leukotriene metabolism, that has previously been hypothesized as a marker of high grade intraepithelial neoplasia (Eisenach et al., 2013).

### POLYPS DEMONSTRATE INCREASED ACTIVITY OF WNT SIGNALING TRANSCRIPTION FACTORS TCF AND LEF

To identify groups of polyps associated with invasive transformation, we clustered the 34,498 peaks that were significantly differential compared to the nearest normal cell type in at least two samples into 10 k-means clusters (Figure 4H), revealing six peak clusters that become more accessible and four clusters that become less accessible at different stages of the transition to cancer. To identify TFs driving chromatin accessibility changes in the transition from normal colon to CRC, we next computed hypergeometric enrichment of motifs in each cluster of peaks from Figure 4H (Figure 4I).

TCF and LEF family motifs were enriched in all clusters that became more accessible across the malignancy continuum, consistent with increased WNT signaling in response to loss of APC leading to increased TCF and LEF activity driving many of the increases in chromatin accessibility that occur in the development of CRC. This observation is consistent with the fact that loss of APC leads to accumulation of β-catenin in the nucleus, which interacts with TCF and LEF transcription factors to drive WNT signaling (Cadigan and Waterman, 2012; Korinek et al., 1997; Morin, 1997). Furthermore, we note that this regulatory transformation is gradual across the malignant continuum—new peaks containing TCF and LEF motifs continue to open at all stages of colon cancer development, as does overall accessibility aggregated across TCF and LEF motifs, suggesting that WNT signaling gradually increases throughout this transformation, over and above what is observed in normal stem cell populations.

Cluster 5 peaks, which became more accessible in later stage polyps and CRC, also exhibited enrichments of ASCL2 motifs (Figure 4I). ASCL2 is a master regulatory of intestinal stem cell fate, and induced deletion of ASCL2 leads to loss of LGR5+ intestinal stem cells in mice (van der Flier et al., 2009). Consistent with a linkage between a more stem-like state in polyp epithelium and more advanced malignant continuum scores, the gene expression of ASCL2 gradually increases as polyps approach malignant transformation (Figure 4J), again indicative of a “super stem” like phenotype, wherein master regulators of stem state are even more active than they are in normal stem cells.

### HNF4A DRIVES THE ACCESSIBILITY OF CANCER SPECIFIC CHROMATIN ACCESSIBILITY

Clusters 4 and 6 exhibit large accessibility increases only in CRC samples, and the greatest enrichment for HNF4A motifs (clusters 4 and 6; Figure 4I). Interestingly, HNF4A expression decreases across polyps along the malignancy continuum, but is then greatly over expressed once the transformation to CRC occurs. These observations may reconcile previous seemingly contradictory observations regarding the role of the transcription factor HNF4A in the development of CRC. On one hand, conditional knockout of HNF4A in adult mice leads to increased proliferation in the crypts and increased WNT signaling, suggesting that HNF4A may be a tumor suppressor gene (Cattin et al., 2009). Similar to this finding, we observe that motifs of HNF4A are most accessible and that HNF4A is highest expressed in mature epithelial cells in normal colon, suggesting that HNF4A is at least associated with less cell proliferation (Figures S4F and S4G). However, other studies have shown that knockdown of HNF4A in CRC cell lines inhibits growth (Schwartz et al., 2009) and that HNF4A is overexpressed in CRC (Cancer Genome Atlas Network, 2012). In our data, HNF4A is gradually lost in polyps, likely leading to an increase in WNT signaling and proliferation, which is consistent with the finding of increased proliferation when HNF4A is overexpressed in normal colon. However, when the transformation to carcinoma occurs, HNF4A becomes overexpressed and begins to drive accessibility of cancer-specific peaks.

### KLF MOTIFS ARE ENRICHED IN PEAKS LOST DURING MALIGNANT TRANSFORMATION

We next examined the four clusters of peaks that became less accessible along the malignancy continuum. Among these clusters were groups of peaks that are less accessible in nearly all polyps (cluster 7) and groups of peaks that only become less accessible in CRC (cluster 10). Motifs enriched in these clusters included HOX family motifs, KLF motifs, and GATA motifs, amongst others. Enrichment of KLF factors is notable as some KLF factors, such as KLF4, have been shown to be tumor suppressors in CRC (Wei et al., 2006). Indeed, loss of accessibility at peaks containing KLF motifs is an early event in the malignancy continuum (cluster 10; Figure 4I).

We observe that many KLF factors, such as *KLF3*, *KLF4*, *KLF5*, *KLF9*, and *KLF12* decrease in expression along the malignancy continuum (Figure S5A). In normal colon differentiation, some KLF factors contribute to maintaining a proliferative state, while others are associated with epithelial differentiation (Kim et al., 2017). We find that these same KLF factors that decrease in expression in stem cells along the malignancy continuum are less expressed in normal stem cells relative to more mature epithelial cells, suggesting that these factors may drive differentiation in normal colon (Figure S5B). As a result, loss of these factors in stem-like cells along the malignancy continuum may prevent differentiation of stem-like cells in polyps and CRC. *KLF7* was an exception to this trend, as expression of *KLF7* increased in stem-like cells along the malignancy continuum (Figure S5A). However, unlike the other factors, *KLF7* was more highly expressed in normal colon stem and cycling cells (Figure S5B) compared with normal enterocytes. This indicates that, unlike other KLF factors, overexpression of *KLF7* in polyps and CRC may also prevent differentiation of stem-like cells.

### REMODELING OF CELLULAR COMPOSITION ALONG THE TRANSITION TO CANCER SHOWS PROGRESSIVE INCREASE IN STEM CELLS, REGULATORY T-CELLS, AND preCAFs

We next asked how the cellular composition of polyps and CRCs change along the continuous malignant transformation. We calculated the fractional contributions of each cell type to each sample as a function of position in the malignancy continuum, and found some cell types were highly correlated with progression along the malignancy continuum. For example, the fraction of stem cells within a sample gradually increases throughout malignant transformation (Figure 5A). Similarly, the number of mature enterocytes decreases as polyps transform to carcinomas (Figure 5B). In the secretory compartment, which primarily consists of immature and mature goblet cells, we observe a fractional increase in immature goblet cells in many of the polyps. In carcinomas we see a pervasive lack of differentiation into the secretory lineage, effectively eliminating immature and mature goblet cells (Figure 5C,5D).

**Figure 5:**
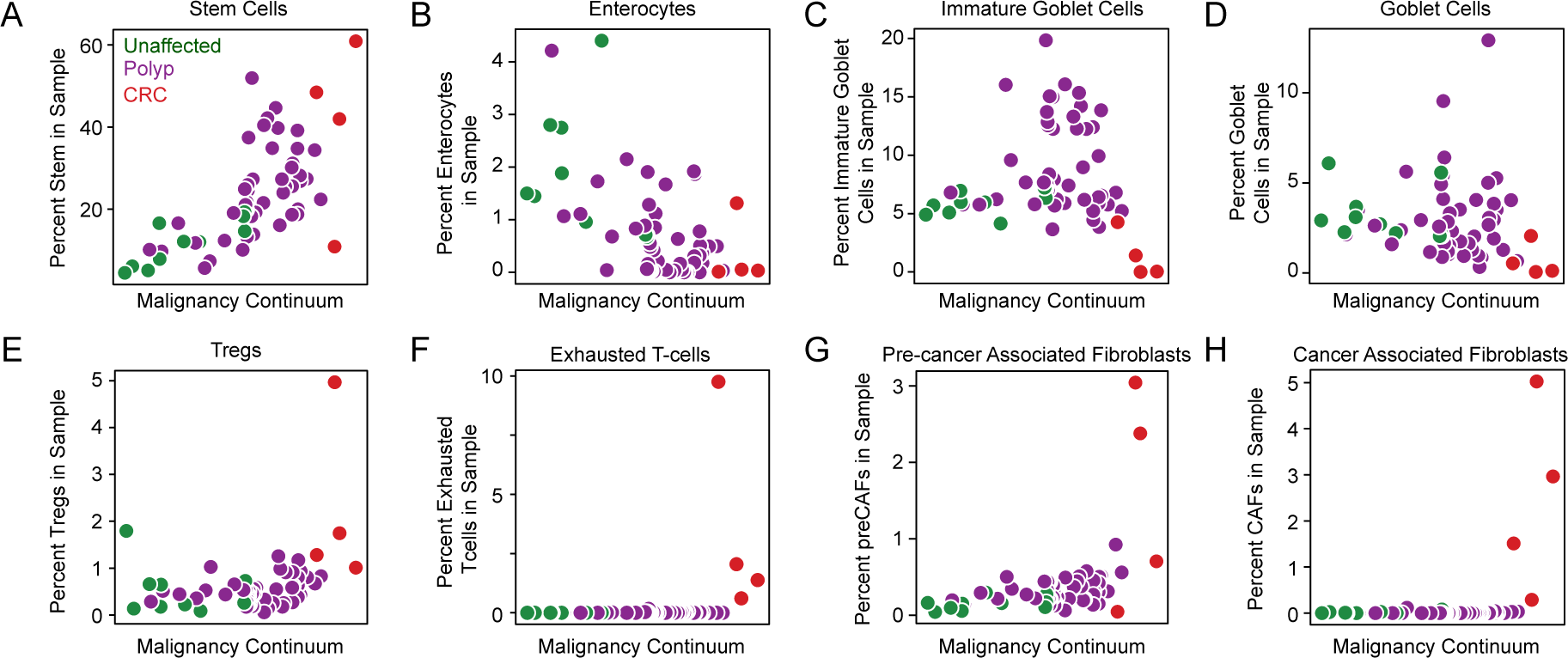
Dynamics of Cell-type Representation in Malignant Transformation. (A–H) Fraction of cell type in each sample plotted against position of the sample in the malignancy continuum defined in Figure 4D for stem-like cells (A), enterocytes (B), immature goblet cells (C), goblet cells (D), Tregs (E), Exhausted T-cells (F), preCAFs (G), and CAFs (H). Samples are colored based on if they are derived from unaffected tissues, polyps, or CRCs.

Outside the epithelial compartment, we also observe dynamic changes in cellular composition that occur progressively during the transformation from unaffected to polyp to carcinoma. Within the immune compartment, we found increasingly large populations of Tregs in the more malignant polyps and CRCs, but exhausted T-cells only in CRCs (Figure 5E,5F). Tregs are known to suppress the antitumor immune response and are typically present at high levels in the tumor microenvironment (Plitas and Rudensky, 2020). The gradual increase in Tregs may be a mechanism of immune evasion in precancerous polpys. Within the stromal compartment, the fraction of preCAFs gradually increases, while CAFs only appear in CRCs (Figure 5G,5H). This observation suggests that preCAFs may support a microenvironment necessary for invasive transformation, and defines a transformation of non-cancerous stromal cells similar to that of the stem-like transformation of cancer cells.

### UPREGULATED SIGNALING RECEPTORS ALONG THE MALIGNANCY CONTINUUM NOMINATES STROMAL-EPITHELIAL INTERACTIONS CONTRIBUTING TO CANCER DEVELOPMENT

To further investigate possible interactions between stromal and epithelial cells, we examined if stem-like epithelial cells in polyps and CRCs expressed receptors that are targets of ligands expressed by CAFs. To do this we first computed potential ligands that were differentially upregulated in CAFs relative to other stromal cell types. We then used the phantom5 database to generate a list of possible receptors for the upregulated ligands. We tested if the differential expression of these receptors in stem-like cells was highly correlated with position along the malignancy continuum. Among the receptors that were most correlated with position along the malignancy continuum were the syndecan family proteins, SDC1 and SDC4 (Figure S4D). Syndecan proteins function as coreceptors for growth factors, matrix proteins, cytokines, and chemokines and have been suggested to both promote or suppress cancer (Sayyad et al., 2019). One possible mechanism by which these proteins could contribute to cancer formation is through signals from the developing tumor microenvironment. We found that a number of genes that are upregulated in CAFs are known to interact with syndicans. For example, SDC4 interacts with ADAM12 (Thodeti et al., 2003), which was previously identified in tumor-associated stroma (Peduto et al., 2006), and is highly expressed in CAFs in our dataset (Figure S4E). Thrombospondin proteins also bind syndicans (Barbouri et al., 2014), and are highly expressed in CAFs.

Another transmembrane receptor that gradually increased in expression along the malignancy continuum was RPSA (Figure S4D), which acts as a cell surface receptor for lamins and is thought to facilitate invasion and metastasis by altering cancer cell interactions with the extracellular matrix (Ménard et al., 1998). We also observed high expression of lamins in CAFs (Figure S4E), which may provide increased lamins to bind RPSA receptors on cancer stem cells. Together, these analyses nominate pairs of ligands and receptors that become increasingly expressed along the malignancy continuum by CAFs and stem-like cells respectively, suggesting mechanisms by which CAFs may work with precancerous cells to contribute to tumorigenesis.

### CHROMATIN ACCESSIBILITY CHANGES ALONG THE MALIGNANCY CONTINUUM ARE STRONGLY ANTI-CORRELATED WITH METHYLATION CHANGES IN CRC

Aberrant DNA methylation is a primary mechanism of tumorigenesis in CRC (Ashktorab and Brim, 2014; Hinoue et al., 2012; Lengauer et al., 1997), but the timing and extent to which methylation changes drive changes in chromatin accessibility prior to and during malignant transformation is not known. To address this, we obtained TCGA DNA methylation data (Illumina 450k array) collected from CRCs and normal samples (Cancer Genome Atlas Network, 2012) and identified probes that were differentially methylated between normal and CRC samples (FigureS6D, Methods). Next, we identified chromatin accessibility peaks from the epithelial cells that overlapped a 450K probe, leaving ∼89,000 peaks overlapping 180,000 probes. For these ∼89,000 peaks, we determined how many overlapped at least 1 hypermethylated site, at least 1 hypomethylated site, or no differentially methylated sites. We then divided the peaks into groups based on whether they were members of significantly upregulated or significantly downregulated clusters of peaks identified in Figure 4H.

For peaks overlapping hypomethylated probes, approximately one third (511) were significantly more accessible in at least 2 samples in our dataset, while < 0.5% (5) became significantly less accessible (Figure 6A; p ≤ 0.05, |Log_2_FC|≥1.5). We saw similar correspondence for peaks overlapping hypermethylated probes, with approximately one quarter (758) becoming significantly less accessible in at least 2 samples in our dataset, while < 0.5% (11) of peaks becoming more accessible (p ≤ 0.05, |Log_2_FC|≥1.5). Therefore, hypermethylation and hypomethylation in CRC nearly perfectly predicts that accessibility at that site will either decrease or increase (respectively), or remain unchanged. When we investigate peaks that do not meet our significance threshold in at least 2 samples, we still observe significantly less aggregate accessibility within peaks overlapping hypermethylated probes (Figure 6B–Non-diff, hypermethylated; 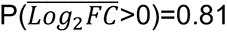, sign test p<10^-50^)) and more accessibility when they overlap hypomethylated probes (Figure 6B–Non-diff, hypomethylated; 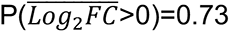, sign test p<10^-50^)). However, we also observe 79.9% (2,189) of significantly more accessible and 75.2% (2,356) of significantly less accessible peaks that overlap probes that are not differentially methylated, implying that a substantial fraction of changes in chromatin accessibility are likely not driven by methylation.

**Figure 6:**
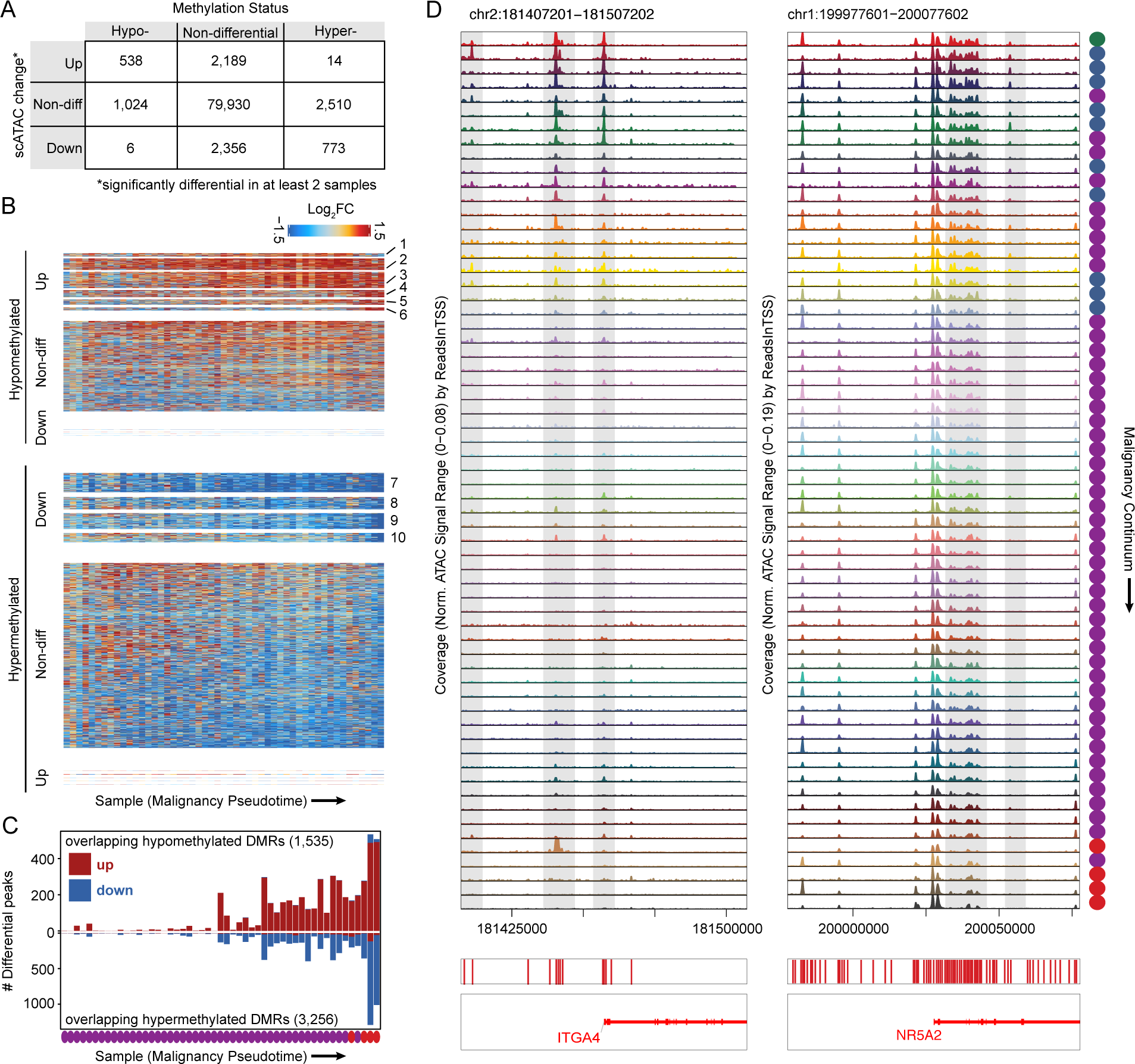
Integration of single-cell colon data with CRC methylation data reveals CRC dmrs with early changes in chromatin accessibility. (A) Table relating the change in accessibility for peaks to the methylation status of illumina 450K methylation probes they overlap. Peaks classified as up were members of clusters 1–6 in Figure 4H and peaks classified as down were members of clusters 7–10 in Figure 4H. (B) Heatmaps of peaks overlapping hypomethylated (top) and hypermethylated (bottom) 450K probes in CRC. The heatmaps are split into peaks from more accessible and less accessible groups defined in Figure 4H and peaks not included in Figure 4H. (C) Number of significantly differential peaks overlapping hypomethylated or hypermethylated 450K probes for each sample. The total number of peaks overlapping hypermethylated and hypomethylated probes is listed in each plot. (D) Accessibility tracks around *ITGA4* and *NR5A2*, which are hypermethylated in CRC. Tracks are ordered by position of the corresponding sample in the malignancy continuum defined in Figure 4.

To identify where in the transition from normal colon to CRC these chromatin changes associated with methylation changes generally occur, we plotted the number of differential peaks overlapping hypermethylated and hypomethylated probes across the malignancy continuum (Figure 6C). Very few of these peaks are differentially accessible in early stage polyps, but substantially more are differentially accessible in late stage polyps and CRC. This observation suggests that changes in chromatin accessibility that occur in regions that are ultimately differentially methylated in CRC accumulate along the transition from normal to cancer.

Among the peaks that both overlap hypermethylated DMRs in CRC and also become less accessible in polyps are a number of previously reported cancer-specific hypermethylated loci (Barault et al., 2018). For example, the promoter region and multiple distal regulatory elements near the *ITGA4* gene are accessible in normal colon, unaffected FAP colon, and very early-stage polyps, but become closed early in the progression to CRC and remain closed even in low-grade polyps (Figure 6D). The gene with the most nearby differential peaks overlapping hypermethylated probes in our dataset was *NR5A2*. Multiple peaks near this gene become less accessible along the malignancy continuum (Figure 6D) and expression of *NR5A2* also gradually decreases along the malignancy continuum (Figure S3D). NR5A2 is a nuclear receptor that has been linked to a wide range of functions including inflammation and cell proliferation. NR5A2 is thought to reduce gut inflammation and increase cell proliferation, which are competing functions in the setting of tumorigenesis (Fernandez-Marcos et al., 2011). We observe downregulation of NR5A2 across methylation, accessibility, and gene expression suggesting that the pro-inflammatory state that may be triggered by the loss of NR5A2 is more important for tumorigenesis than this loss might have on cell proliferation.

Hypermethylated DNA regions in CRC have also been incorporated into commercial tests screening for CRC. For example, Cologuard includes the hypermethylation of the promoter regions of *BMP3* and *NDRG4* (Imperiale et al., 2014). We examined how accessibility changes around BMP3 across our malignancy continuum and found multiple peaks that become inaccessible in the middle of the continuum (Figure S6A). This likely represents the point that these regions become hypermethylated and helps segregate which of these polyps would test positive or negative for this biomarker. We observe many regions with a similar behavior: sharp increases or decreases in accessibility at a specific point along the malignancy continuum. We speculate that testing for accessibility, or likely methylation, at these loci would likely enable not just calling polyps or CRCs but also staging the polyp along the malignancy continuum. This approach also identifies methylation markers/loci (e.g., *GRASP*, *CIDEB*) specific for malignant transformation in CRC (Figures S6B, S6C).

## DISCUSSION

Strategies to identify cancers in a premalignant stage, where interventions might be highly efficacious, promise tractable means to prevent cancer deaths. However, most previous work profiling genetic, epigenetic, and transcriptomic changes that occur in malignancy has focused on advanced tumors rather than premalignant lesions. Our single-cell atlas of colon cancer tumorigenesis fills this gap by identifying key changes in chromatin accessibility, gene expression, and tissue composition that occur along this transformation, and provides a wealth of potential targets for prevention, diagnosis, and treatment of malignancy.

Unlike bulk assays, which convolve changes in cell composition with changes in cell state, our single-cell approach identified both dramatic changes in cell composition and cell state that occur during the malignant transformation. Analysis of both the composition and cell state of precancerous epithelial cells revealed that an increasing fraction of epithelial cells occupy a stem-like state as polyps approach malignancy, but these cells also exhibit underlying dysfunctional epigenetic and gene expression programs distinct from normal stem cells. We also identify compositional and cell state changes of non-cancerous cells, including fibroblasts and immune cells, which may be influencing the tumor microenvironment to drive cancer progression.

Gene expression changes along the malignancy continuum implicate mechanisms of cancer initiation and nominate diagnostic and therapeutic targets. We find that expression of *GPX2*, a glutathione peroxidase, is gradually upregulated across the transformation from normal tissue to malignancy. Even in premalignant tissues, *GPX2* is upregulated, suggesting that its role in reducing the oxidative environment may be needed for progression along this continuum. As a result, we speculate that expression of *GPX2* could serve as a marker for the degree of polyp malignancy, and that inhibitors of GPX enzymes—such as tiopronin (Hall et al., 2014)—may be relevant treatment strategies for CRC or premalignant lesions.

We identified a number of transcription factors that appear to drive chromatin accessibility changes as polyps transition to malignancy. In stem-like cells obtained from polyps, we observe that many TCF and LEF motifs inaccessible in normal intestinal stem cells become accessible, suggesting that WNT signaling increases along the malignant continuum over and above that of normal stem cells.

The final step in cancer formation is malignant transformation, and we identify a number of changes in chromatin accessibility, gene expression, and tissue composition associated with this transformation (Figures 4, S3, 5, and S4). Our data identify HNF4A as a key regulator of malignant transformation—as both the expression of this gene becomes upregulated and chromatin regions containing HNF4A motifs become accessible only after the transformation to CRC. In normal colon, HNF4A motifs are more accessible in mature enterocytes than stem cells, but in colon cancer stem cells HNF4A is upregulated and begins driving chromatin accessibility changes not observed in normal stem cells.

Previous work has identified a number of epigenetic factors associated with colorectal cancers, and the methylation state markers are at the core of widely adopted screening tests for CRC (Imperiale et al., 2014). However, little is known regarding if and when the methylation state used in these tests can be observed in premalignant lesions. We find these regions lose their accessibility in the middle of the malignancy continuum, and behave similarly to many thousands of other differentially accessible sites. Future efforts will be needed to untangle the relative timing of chromatin accessibility changes vs. methylation changes, along this continuum. Because we observe that hypermethylated regions are strongly decreased in their accessibility and hypomethylated regions are strongly enriched for increased accessibility, our dataset can be used to stratify differentially methylated regions expected to be present early in the continuum of malignant transformation, those expected to be present later in the continuum, and those expected to be present only in CRC, opening the possibility for a stage-specific methylation-based screening.

Currently, clinical guidelines for the timing and frequency of follow-up endoscopic screening after polypectomy depends on the size of the polyps removed and whether they are classified as low-grade or high-grade dysplasia (Gupta et al., 2020). This work presents a strategy by which to order premalignant polyps by their degree of malignancy, which could be useful for staging polyps and assessing clinical risk. Perhaps the most straightforward approach to this type of staging is to use the fraction of stem-like cells (through IHC labeling of cell-type specific markers such as LGR5) as a proxy for position along the malignancy pseudotime trajectory (see Figures 4, S3, and S4). We also observe that the fraction of Tregs in a sample strongly correlates with the position on the malignancy continuum, suggesting that the fraction of Tregs in a polyp may also be a fruitful proxy of malignancy. However, we do not currently know if outcomes would improve if more information was used to define colonoscopy follow-up time, and assessment of this question will require substantial clinical investigation.

This work demonstrates that adenomatous polyps traverse a strikingly consistent epigenetic and transcriptional trajectory as they progress to CRC. These results lead us to question if a similar relatively uniform molecular phenotypic trajectory is common to other premalignant lesions that precede other cancers, or if other pre-cancers might traverse multiple diverse paths on the way to malignant transformation. Further, we might also ask if colorectal cancers continue to follow a consistent pathway to metastasis or if multiple distinct pathways are taken once malignant transformation occurs. We anticipate that similar single-cell integrative methods can be deployed to answer these questions.

## ACKNOWLEDGEMENTS

This work is supported by NIH Grants U2C CA233311, S10OD025212 and P30DK116074. W.R.B. was supported in part by the Stanford MSTP training grant T32GM007365 and the SIGF affiliated with ChEM-H. S.A.N. was supported in part by the Stanford Graduate Fellowship in Science and Engineering. Yiing Lin, MD, PhD of Washington University in St. Louis provided the B001 and B004 tissue samples.

## AUTHOR CONTRIBUTIONS

Conceptualization, W.R.B., S.A.N., M.P.S., W.J.G.; Investigation-Single-cell Experiments, S.A.N., R.C., D.C.C., W.R.B.; Investigation-Pathology Reads, T.L., J.S.; Formal Analysis, W.R.B.; Resources-Sample Collection, R.L., M.M., A.M.H.,U.L., E.D.E., R.C., S.A.N., W.R.B.; Resources-Sequencing, H.C.; Writing-Original Draft, W.R.B., S.A.N., W.J.G., Writing-Review and Editing, All Authors; Funding Acquisition, A.K., C.C., E.D.E, J.M.F., M.P.S., W.J.G.; Supervision, E.D.E., A.K., J.M.F., C.C., M.P.S., W.J.G.

## SUPPLEMENTAL TABLES

Supplemental Table S1: Metadata for patients included in the study, including genetic disease status, age, sex, and race/ethnicity.

Supplemental Table S2: Metadata for individual samples including microscopic pathology reads.

---

**Figure S1:**
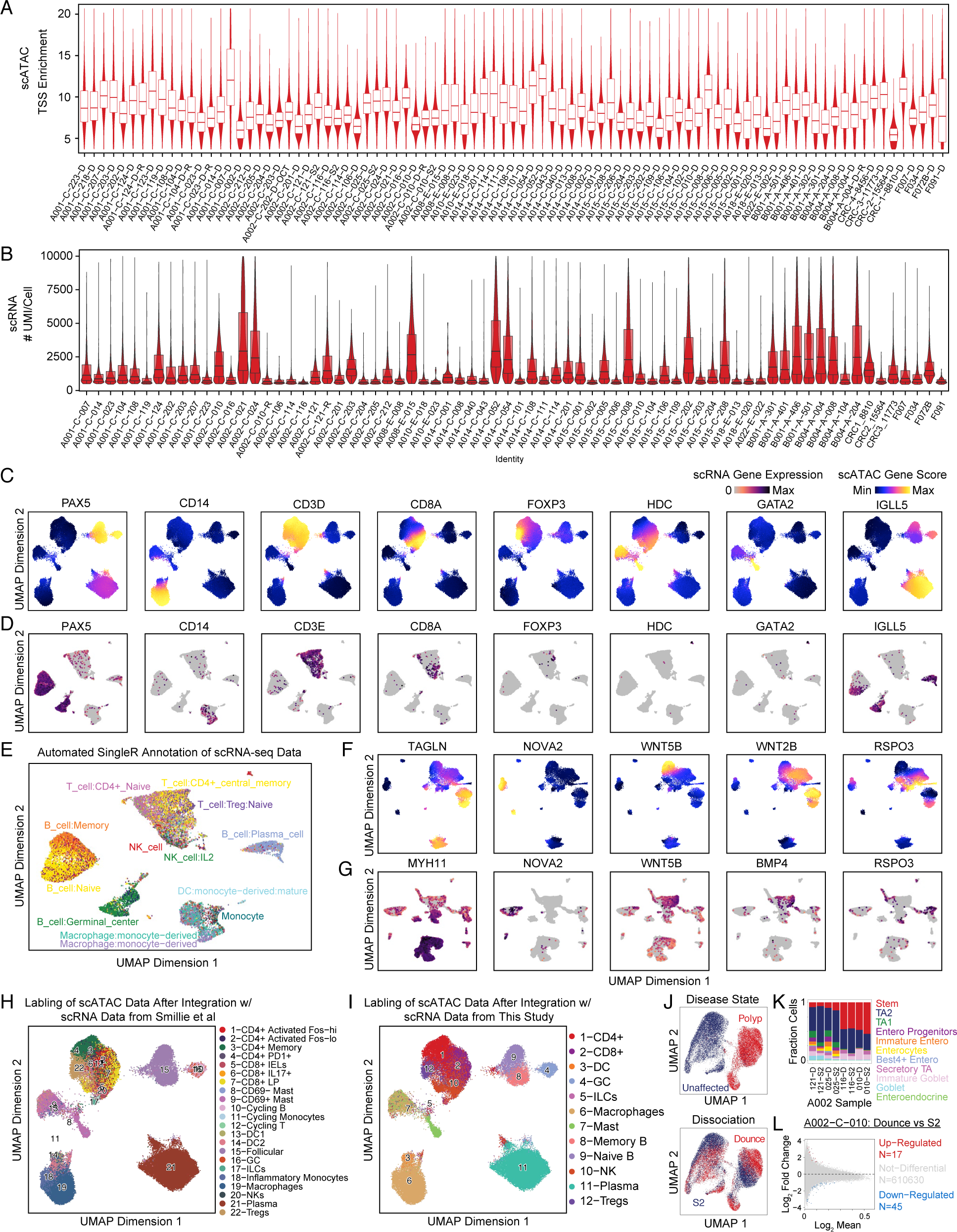
Quality control and annotation of single-cell datasets. (A) Violin plots of TSS-enrichments for all scATAC cells from each sample. Samples are labeled by patient (e.g. A001, A002, etc), source (C=Colectomy, E=Colonoscopy, A=Autopsy, T=Tissue Bank), dissociation (D=dounce, S2 = S2 singulator). Replicates performed on additional sections of the same polyp are indicated with a R. (B) Violin plots of the number of UMIs sequenced for cells from each sample. Samples are labeled the same as in S1A, except all tissues were denounced so the dissociation method is not included. (C) UMAP projection of scATAC immune cells colored by gene activity scores reflecting accessibility within and around immune marker genes. (D) UMAP projection of snRNA immune cells colored by expression of immune marker genes in each cell. (E) UMAP projection of snRNA immune cells colored by automated labeling of snRNA immune cells with SingleR. (F) UMAP projection of scATAC stromal cells colored by gene activity scores of stromal marker genes. (G) UMAP projection of snRNA stromal cells colored by expression of marker genes. (H, I) UMAP projection of scATAC immune cells where cells are labeled by the nearest snRNA cell from (H) Smillie et al or (I) this study after integrating the respective datasets with CCA. (J) UMAP projections of four scATAC samples with nuclei isolated with both douncing and the S2 Singulator, colored by disease state (top) and dissociation method (bottom). (K) Fraction of epithelial cells of each cell type for the 4 samples where nuclei were isolated with douncing and the S2 singulator. (L) Differential peaks between scATAC stem cells isolated from two sections of the same polyp that were processed with either the S2 singulator or douncing.

**Figure S2:**
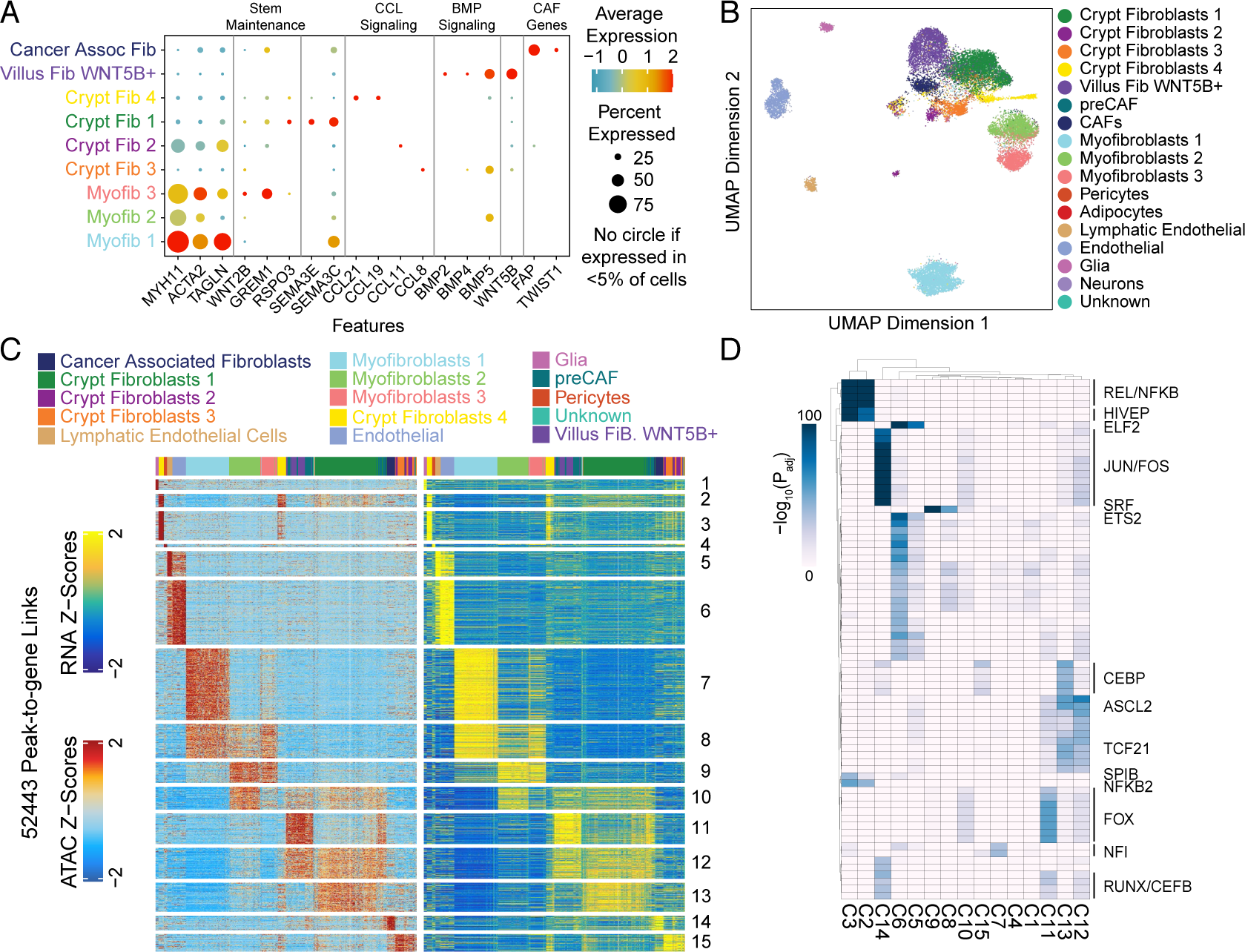
Cell-type specific expression and RNA-ATAC integration of stromal cells. (A) Dotplot representation of RNA expression of myofibroblast, stem maintenance, SEMA, CCL, BMP, and CAF genes by cells in different fibroblast subtypes. (B) Labeling of scATAC cells by aligning scATAC and snRNA data with CCA and labeling scATAC cells based on nearest snRNA cells. (C) Peak-to-gene linkages between scATAC and snRNA stromal cells. Rows in the left heatmap represent peaks and are colored by accessibility while rows in the right heatmap represent genes and are colored by expression. (D) Hypergeometric enrichment of motifs in clusters of peaks from S2C.

**Figure S3:**
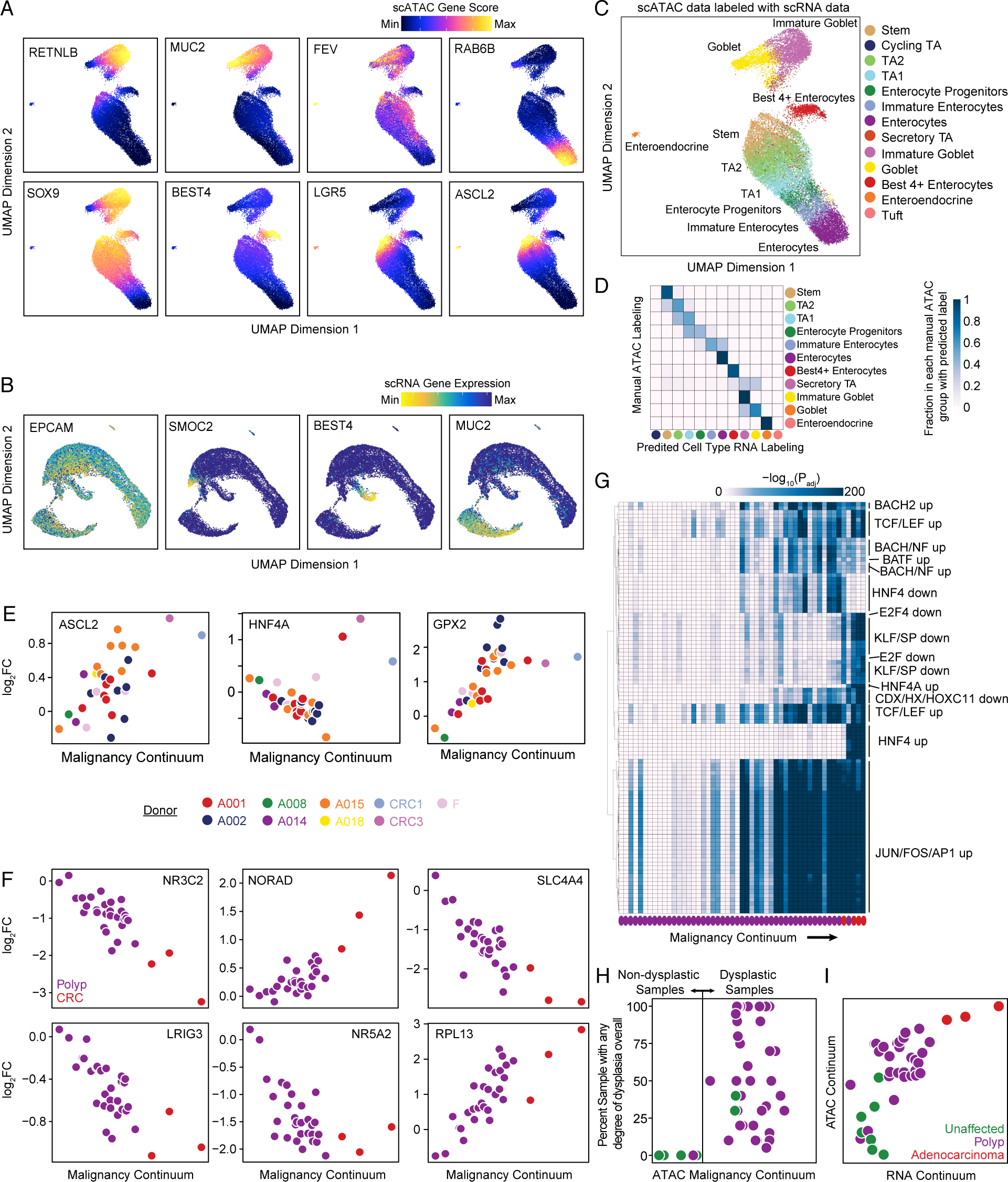
Characterization of normal colon epithelium and identification of changes along the malignancy continuum. (A) UMAP projection of normal colon epithelial cells colored by scATAC gene activity scores of the epithelial marker genes *RETNLB* (immature goblet), *MUC2* (goblet), *FEV* (enteroendocrine), *RAB6B* (enterocyte), *SOX9* (stem), *BEST4* (Best4+ enterocyte), *LGR5* (stem), and *ASCL2* (stem). (B) UMAP projection of normal colon epithelial cells colored by expression of marker genes *EPCAM* (general epithelial), *SMOC2* (stem), *BEST4*, and *MUC2*. (C) Labeling of scATAC epithelial cells by nearest snRNA cells following integration of the datasets with CCA. (D) Confusion matrix comparing annotation of scATAC cells using marker genes and labeling of scATAC cells with the nearest snRNA cell following integration of scATAC and snRNA datasets. (E) Log_2_FC in expression of *ASCL2*, *HNF4A*, and *GPX2* in stem-like cells from each sample relative to stem-like cells in unaffected samples plotted against the malignancy continuum defined in 4D. Samples are colored based on the patient the sample was collected from. (F) Log_2_FC in expression of *NR3C2*, *NORAD*, *SLC4A4*, *LRIG3*, *NR5A2*, and *RPL13* as a function of malignancy continuum. Samples are colored based on if they are derived from polyps or CRCs. (G) Enrichment of motifs in differential peaks identified for stem-like cells for each sample relative to stem-like cells in unaffected samples. Up after the motif name indicates that the motif is enriched in peaks that become more accessible in the sample and down indicates that the motif is enriched in peaks that become less accessible. While the analysis in Figure 4 focuses on groups of peaks that change in CRC, this analysis highlights peaks that change within individual samples to test if specific samples are associated with gain or loss of specific peaks. (H) Relationship between the malignancy continuum and percent of sample with any degree of dysplasia as determined by microscopic pathology. Samples are colored based on gross classification as a polyp (purple) or unaffected (green) tissue. Note that some samples classified as unaffected had dysplasia while some samples classified as polyps did not have dysplasia. (I) Relationship between the malignancy continuums defined from the scATAC and scRNA datasets. Samples are colored based on gross classification as unaffected, polyp, or CRC.

**Figure S4:**
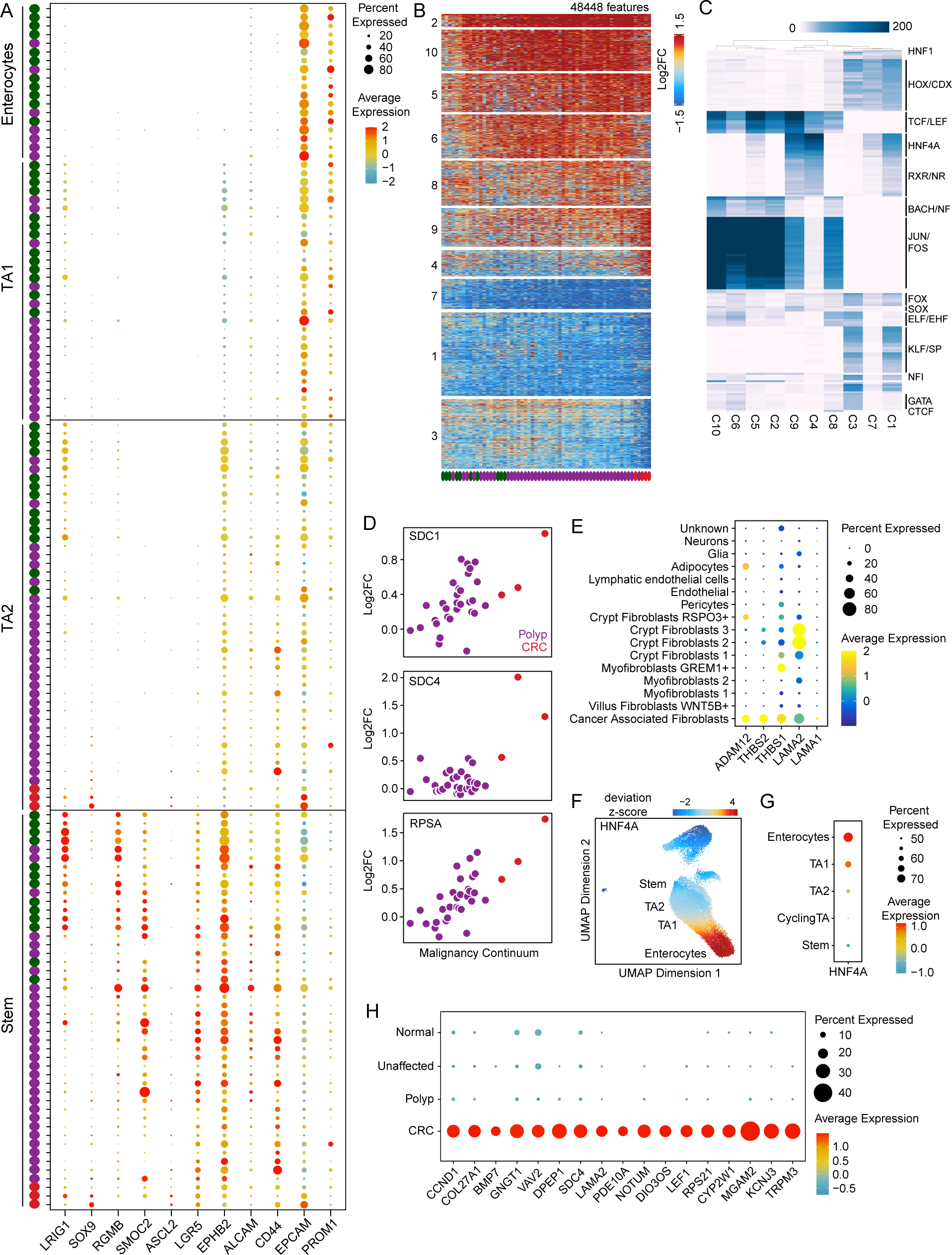
Epigenetic and transcriptomic changes along the malignancy continuum. (A) Expression of intestinal stem cell and colon cancer stem cell marker genes in stem cells, TA2 cells, TA1 cells, and Enterocytes by sample. Samples are ordered by the malignancy continuum defined in Figure 4D. (B) Heatmap of all peaks that were significantly differentially accessible in ≥2 samples between stem cells from a given sample and normal colon stem cells. Samples are ordered along the x-axis by the malignancy continuum defined in 4D. Peaks are kmeans clustered into 10 groups. (C) Hypergeometric enrichment of motifs in clusters of peaks defined in S4B. (D) Log_2_FC in expression of *SDC1*, *SDC4*, and *RPSA* along the malignancy continuum. Samples are colored based on if they are from polyps (purple) or CRC (red). (E) Dotplot representation of the expression of selected ligands by different stromal cell types. (F) UMAP projection of normal colon epithelial cells colored by motif activity of HNF4A. (G) Dot plot representation of *HNF4A* expression in different normal colon epithelial subtypes. (H) Dot plot representation of genes differentially expressed in CRC relative to polyps.

**Figure S5:**
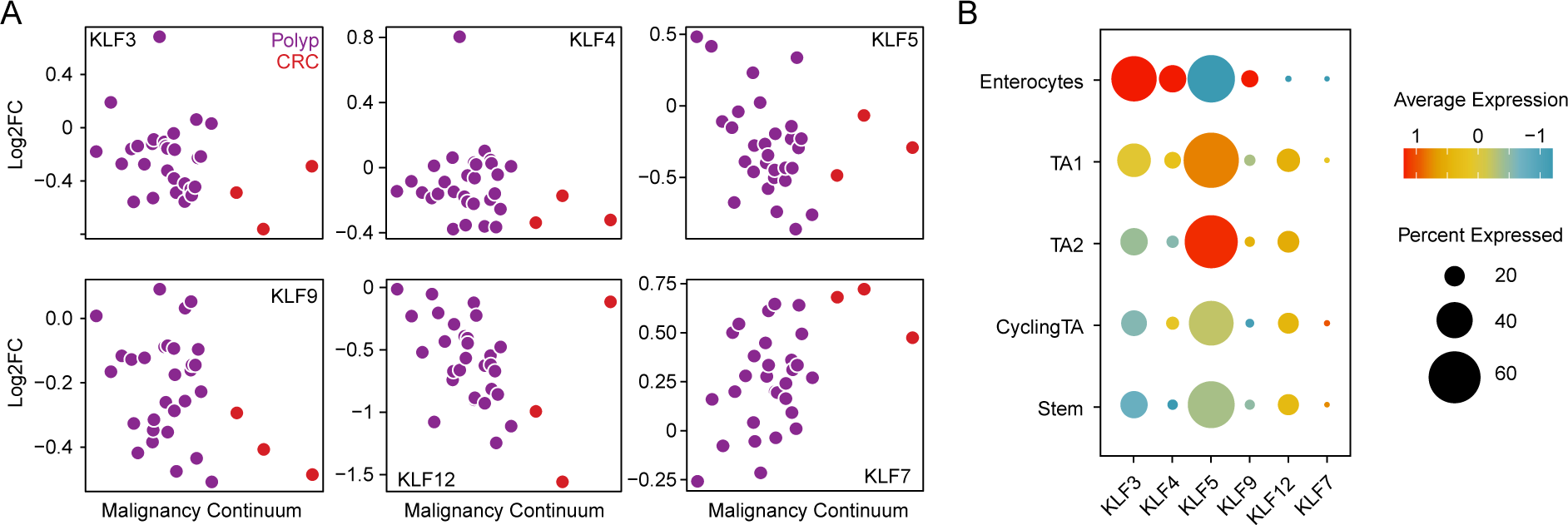
Expression of KLF transcription factors along the malignancy continuum. (A) Log_2_FC in expression of KLF transcription factors relative to unaffected colon as a function of position along the malignancy continuum. Samples are colored based on if they are from polyps (purple) or CRC (red). (B) Dotplot representation of the expression of KLF transcription factors in normal colon epithelial cells.

**Figure S6:**
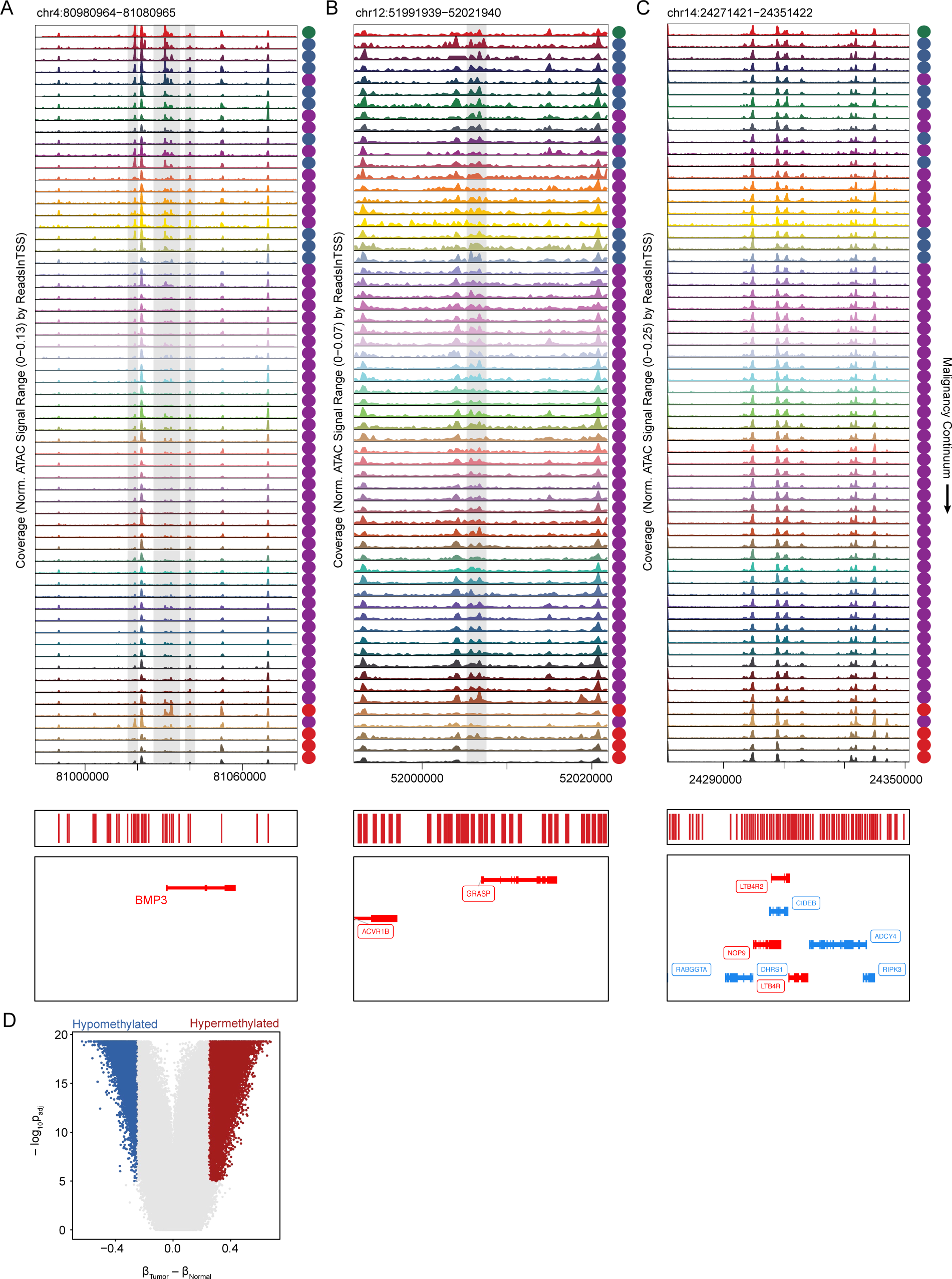
Accessibility changes in regions hypermethylated in CRC. (A–C) Accessibility tracks around *BMP3* (A), *GRASP* (B), and *CIDEB* (C), which are hypermethylated in CRC. (D) Adjusted p-value and mean difference in β-value cutoffs used to determine differentially methylated probes.

## METHODS

### Experimental Methods

#### Description of FAP donors

We collected samples from the following groups of patients: FAP (8 patients), routine colonoscopy screening (1 patient), sporadic CRC (4 patients), and healthy controls (2 patients). FAP tissue was collected at the time of partial or full colectomies for 4 patients and during screening colonoscopies for 4 patients. From non-FAP patients, 1 sporadic polyp was obtained during a standard colonoscopy as part of routine screening, 4 sporadic CRCs were obtained from the Stanford Tissue Bank, and 8 normal tissue samples were collected from brain-dead organ donors under consent.

Patient-matched normal colon mucosa, polyps, and adenocarcinomas were flash frozen in liquid nitrogen at time of collection and stored at -80°C. 1 polyp (A002-C-202) and 1 adenocarcinoma (A001-C-007) were embedded in optimal cutting temperature compound (OCT) prior to storage at -80°C. Polyps were scored by a board-certified pathologist for presence of dysplasia, including low or high grade with corresponding percentages. For adenocarcinoma samples, staging was performed. A small number of polyps (7 polyps) were exhausted for molecular assays so pathology reads were not obtained.

#### Tissue Dissociation and Nuclei Isolation

All protocols used to generate snRNA-seq data on the 10x Chromium platform, including sample prep, library prep, instrument, and sequencing settings, can be found on the 10x Genomics website at: https://support.10xgenomics.com/single-cell-gene-expression. The isolation of nuclei was accomplished using the OmniATAC protocol (Corces & Granja et. al. Science 2018). Subsequent downstream GEM barcoding, cDNA construction, and gene expression library construction were performed according to the Chromium Next GEM Single Cell 3’ Reagent Kits v3.1 (10x Genomics, 1000121) and Chromium Next GEM Single Cell ATAC Library & Gel Bead Kit v1.1 (10x Genomics, 1000175). Dissociation of nuclei was carried out entirely on wet ice. 40-60mg of flash-frozen tissue was placed into 2ml dounce tissue grinders containing 1ml HB (Lysis) Buffer (1.0341x HB Stable solution, 1M DTT, 500 mM Spermidine, 150mM Spermine, 10% NP40, complete Protease Inhibitor, Ribolock), gently triturated, then allowed to thaw for 5 minutes in the solution. After 5 minutes, tissue was dounced 10 times with pestle A and 20 times with pestle B, or until there was no resistance from either pestle. Sample was filtered through a 40um cell strainer (Falcon; 352340) and resulting homogenate transferred to a pre-chilled 2ml LoBind tube. Samples were spun in a 4°C fixed angle centrifuge for 5 minutes at 350 RCF to pellet nuclei. After spinning, all but 50ul of supernatant was removed. 350ul HB was added to the nuclei pellet for a total volume of 400ul. Nuclei were gently resuspended with a wide bore pipet. One volume of 50% Iodixanol (60% OptiPrep [Sigma Aldrich; D1556], Diluent Buffer [2M KCl, 1M MgCl2, 0.75M Tricine-KOH pH 7.8], Water) was added and gently triturated. Next, 600ul of 30% Iodixanol was carefully layered under the 25% mixture. Finally, 600ul of 40% Iodixanol was layered under the 30% mixture. Sample was then spun in a 4°C swinging bucket centrifuge for 20 min at 3,000 RCF. Upon completion of the spin, a band of nuclei was visible. Supernatant was aspirated down to within 200-300ul of the nuclei band. Nuclei band was then collected at 200ul and transferred to a fresh 1.5ml tube. Sample was diluted with one volume (200ul) Resuspension Buffer (1x PBS, 1% BSA, 0.2u/uL Ribolock). Nuclei concentration was determined using the Countess II FL Automated Cell Counter (ThermoFisher; AMQAF1000). Average time from dissociation to generating barcoded GEMs for snRNA-seq was 30-45 minutes.

In addition to isolating nuclei with the omni-ATAC dounce method, we used the Singulator from S2 Genomics to isolate nuclei for 4 samples (2 normal and 2 polyp) from patient A002. The same buffers (lysis, wash, resuspension) were used for both the manual omni-ATAC dounce and S2 Singulator methods. Data generated from nuclei dissociated with the Omni-ATAC protocol and S2 genomics exhibited excellent agreement (Figures S1J–S1L). HB buffer was prepared following the OmniATAC protocol, with the addition of 15ul Ribolock per 1mL HB buffer to help maintain RNA integrity. Approximately 50-70mg tissue in small chunks were placed into the Nuclei Isolation Cartridge. We ran the Extended Nuclei Isolation Singulator program which includes disruption, 5 minute incubation, disruption, filtration (150um and 40um filters) and buffer rinse of the cartridge. Following dissociation, nuclei were spun at 300g for 5 minutes. Supernatant was removed and nuclei were resuspended in Resuspension Buffer (noted above). Nuclei were counted using the Countess Automated Cell Counter (ThermoFisher). The total dissociation time for 4 samples was approximately 1 hour, as one sample can be run at a time on the S2 Singulator. Nuclei were immediately used for single cell sample prep and library construction following the 10x Chromium NextGEM Single Cell 3’ V3.1 protocol and Single Cell ATAC v.1.1 protocols (10x Genomics).

#### Single-cell assay for transposase-accessible chromatin using sequencing (scATAC-seq)

scATAC-seq targeting 9,000 cells per sample was performed using Chromium Next GEM Single Cell ATAC Library & Gel Bead Kit v1.1 (10x Genomics, 1000175) and Chromium Next GEM Chip H (10x Genomics, 1000161). Each sample library was uniquely barcoded and quantified by qPCR using a PhiX Control v3 (Illumina, FC-110-3001) standard curve. Libraries were then pooled and loaded on a NovaSeq 6000 Illumina sequencer (1.4 pM loading concentration, 50 × 8 × 16 × 49 bp read configuration) and sequenced targeting 25,000 reads per cell on average.

#### Single-nuclear whole transcriptome sequencing (snRNA-seq)

snRNA-seq targeting 9,000 cells per sample was performed using Chromium Next GEM Single Cell 3’ Reagent Kits v3.1 (10x Genomics, 1000121) and Chromium Next GEM Chip G Single Cell Kit (10x Genomics, 1000120). Each sample library was uniquely barcoded and quantified by qPCR using a PhiX Control (Illumina, FC-110-3001) standard curve. Libraries were pooled and loaded onto NovaSeq 6000 Illumina sequencer (Read 1 = 28bp, i7 index=8bp, i5 index=0bp, Read 2=91bp read configuration) and sequenced to 20,000 reads per cell on average.

### Analytical Methods

#### Single-cell assay for transposase-accessible chromatin using sequencing

Initial processing of scATAC-seq data was performed using the Cell Ranger ATAC Pipeline (go.10xgenomics.com/scATAC/cell-ranger-ATAC) by first running cellranger-atac mkfastq to demultiplex the bcl files and then running cellranger-atac count to generate scATAC fragments files. These fragments files were loaded into R (v3.6.1) using the createArrowFiles function in ArchR (Granja et al., 2020). QC metrics were computed for each cell, and cells with TSS enrichments less than 4 were filtered out for all samples. Cells were also filtered based on the number of unique fragments sequenced using a cutoff defined for each sample. As the sequencing depth sometimes differed between samples, it was necessary to assign this fragment cutoff as a sample-specific parameter. The sample-specific cutoffs ranged from 1,500 to 10,000, with the most common cutoff being 3000 fragments/cell.

#### Single-cell ATAC-seq: Doublet removal and initial clustering

After generating arrow files for each sample, an ArchR project containing all samples was created. Doublets were simulated using the ArchR function addDoubletScores with k = 10. Cells with the highest probability of being doublets were removed using the ArchR function filterDoublets with a filterRatio of 1.2. Following initial doublet removal, sample wise quality control statistics including TSS enrichment and fragments per cell were recomputed (Figure S1A). As part of construction of the arrow files, a tile matrix consisting of reads in 500bp tiles was constructed. This tile matrix was used as input to compute an iterativeLSI dimensionality reduction using the addIterativeLSI function in ArchR with a total of 2 iterations, a clustering resolution of 0.2 following the first iteration, 25,000 variable features, 30 dimensions, and sampling 50,000 cells. Following dimensionality reduction, initial clustering was performed using ArchR’s addClusters, which is a wrapper for Seurat’s FindClusters function (Stuart et al., 2019) using a resolution of 1.7 and sampling 50,000 cells for clustering (remaining cells were then grouped into clusters based on the nearest cells included in the clustering). We next ran addUMAP on the IterativeLSI dimensionality reduction with 30 nearest neighbors and a minimum distance of 50.

As part of arrow file creation, ArchR computes gene activity scores for each gene. These gene activity scores are a function of the accessibility within and around a given gene body and can be used as a proxy for gene expression. After initial dimensionality reduction and clustering of all cells in our dataset, we examined gene activity scores of known marker genes for cell types expected to be present in either epithelial, stromal, or immune cells and divided cells from our dataset into three groups (immune, stromal, or epithelial) for downstream analysis.

#### Single-cell ATAC-seq: Analysis of Immune Compartment

After subsetting the dataset to include only immune cells, we repeated dimensionality reduction and clustering. The iterative LSI dimensionality reduction was computed using addIterativeLSI with 2 iterations, 25000 variable features, sampleCellsPre set to NULL, dimensions 1–30, and a clustering resolution of 0.2 for the initial iteration. Clusters were then determined with addClusters in ArchR using the Seurat method, a resolution of 1.7, and nOutlier set to 50. The UMAP dimensionality reduction was then computed using addUMAP with 30 nearest neighbors, a minimum distance of 0.5, and the cosine metric. We next examined gene activity scores of known marker genes and identified 3 small clusters with gene activity scores for marker genes that were not consistent with these clusters consisting of a single high-quality immune cell subtype. As a result, these clusters were thought to be likely doublets, and were removed prior to additional analysis. We note this approach to removing likely doublet clusters is commonly employed for single-cell datasets and a similar approach has recently been applied to remove doubles in a large snRNA-seq dataset on colon cells (Smillie et al., 2019). Following this additional double removal step, the dimensionality reduction and clustering steps were repeated using identical parameters.

Multiple approaches were taken to annotate the scATAC data. First, gene activity scores of known marker genes for different immune populations expected to be present in the scATAC data were examined for the different scATAC clusters. Marker genes included *PAX5*, *MS4A1*, *CD19*, *IGLL5*, and *VPREB3* for B-Cells; *TPSAB1*, *HDC*, *CTSG*, *CMA1*, *KRT1*, *IL1RAPL1*, and *GATA2* for Mast cells; *KLRF1*, *SH2D1B*, and *SH2D1B* for NK cells; *SSR4*, *IGLL5*, *IGLL1*, and *AMPD1* for Plasma cells; *CD14* for Monocytes; *CD3D*, *CD3E*, *CD3G*, *CD8A*, *CD8B*, *TBX21*, *IL7R*, *CD4*, *CD2*, *BATF*, *TNFRSF4*, *FOXP3*, *CTLA4*, and *LAIR2* for T-cells and T-cell subtypes; and *FOLR2*, *FABP3*, and *PLA2G2D* for Macrophages. This approach led to unambiguous identification of most clusters in our dataset. While annotating the clusters, some clusters were labeled with the same annotation if they consisted of cells of the same subtype. We next integrated our scATAC data with multiple snRNA-seq datasets using ArchR’s addGeneIntegrationMatrix function, and then labeled scATAC cells based on the nearest snRNA cells. This included large, high-quality scRNA-seq data from cells isolated from normal colon and UC patients (Smillie et al., 2019). This dataset contains slightly different populations of cells (e.g. no exhausted T-cells) so was insufficient to be used for annotation of our data in isolation. However, we observed good overall agreement between the marker gene based annotations of our scATAC data and the annotations obtained when labeling our scATAC cells with the nearest scRNA cell in the Smillie et al dataset. We also integrated our data with the labeled snRNA data produced in this study, which also produced good agreement with our initial manual labeling. Ultimately, our final annotations are the result of both initial annotation with known marker genes and refinement and validation of our clusters by integrating our scATAC data with multiple scRNA-seq datasets and labeling our scATAC cells with the nearest scRNA cells from these datasets.

We next selected all T-cells present in our scATAC dataset to further explore differences in the T-cell subtypes. Using only the T-cells in our dataset, we ran addIterativeLSI with the same parameters as the full immune data. The cells were clustered using addClusters with a resolution of 2.5 and nOutlier of 50. Impute weights were computed for the T-cell subset and imputed gene activity scores of T-cell marker genes were plotted on the T-cell UMAP with the ArchR function plotEmbedding. The scATAC T-cell dataset was integrated with a previously published dataset consisting of T-cells from BCC (Yost et al., 2019) using addGeneIntegrationMatrix as described above.

T-cell specific peaks were called using addReproduciblePeakSet in ArchR, which generates pseudobulk replicates for user defined groups, calls peaks on these pseudobulk replicates using Macs2 to define a reproducible peak set for each group, and then merges the resulting peak sets for each group into a union peak set of fixed width peaks. For this peak calling step, cells were divided into groups based on T-cell subtype. After calling Macs2, a peak matrix was constructed in ArchR using addPeakMatrix. Annotations of motifs present in the peak set were identified with addMotifAnnotations and the cisbp motif set. Then background peaks were identified with addBgdPeaks and ChromVar deviations were added with addDeviationsMatrix. Imputed deviation z-scores were then plotted on the T-cell UMAP. Differential peaks between different T-cell subtypes were computed with getMarkerFeatures using the wilcoxon test and with bias set to c(“TSSEnrichment”, “log10(nFrags)”). Hypergeometric enrichment of motifs in markerPeaks were computed with peakAnnoEnrichment.

#### Single-cell ATAC-seq: Analysis of Stromal Compartment

To analyze the stromal cells present in our scATAC-seq dataset, we first performed dimensionality reduction and clustering. The iterative LSI dimensionality reduction was computed using addIterativeLSI with 3 iterations, clustering resolutions of 0.1 and 0.2 after the first and second iteration respectively, 20000 variable features, sampleCellsPre set to NULL, and using dimensions 1–30. Clusters were then determined with addClusters using the Seurat method, a resolution of 1.0, and nOutlier set to 50. As was done for the immune cells, we identified marker gene activity scores for each cluster and examined gene activity scores of known marker genes. For the stromal cells, one cluster was identified as a likely doublet cluster and removed. Dimensionality reduction and clustering was then repeated with identical parameters except a resolution of 1.1 was used in the final clustering step. A UMAP dimensionality reduction was then computed using addUMAP with nNeighbors of 35, minDist of 0.5, and the cosine metric.

To assign cell type annotations to the stromal clusters, gene activity scores for known marker genes were examined for different scATAC clusters. The scATAC data was also integrated with snRNA from this study using the ArchR function addGeneIntegrationMatrix, enabling labeling of the scATAC cells based on the nearest snRNA cells. Similar to the immune cells, marker gene activity scores were used for initial annotation while labels from the RNA integration were used to validate and refine the initial annotations to produce the final cell-type annotations.

Following cell type annotation, a stromal cell peak set was generated by first running addGroupCoverages and grouping by CellType and then running addReproduciblePeakSet. To identify markerPeaks for the stromal compartment, getMarkerFeatures was run using the Wilcoxon test method and with TSSEnrichment and log10(nFrags) provided as bias parameters. Marker peaks with FDR <= 0.1 & Log_2_FC >= 0.5 were visualized on the heatmap in Figure 2. Motif annotations using cisbp motifs were then added to the project and enrichment of motifs in marker peaks for each cell type were identified with the ArchR function enrichMotifs and cutoffs of FDR <= 0.1 & Log_2_FC >= 0.5. ChromVAR deviations were then computed using addBgdPeaks and addDeviationsMatrix. Putative peak-to-gene links were identified with the ArchR function addPeak2GeneLinks with default parameters. TFs likely regulating chromatin accessibility were then determined exactly following the ArchR manual for identifying TF Regulators.

The stromal cell trajectory from villus fibroblasts to CAFs was constructed by defining the trajectory as trajectory <-c(“Villus Fibroblasts WNT5B+”, “Inflammatory Fibroblasts”, “Cancer Associated Fibroblasts”) and then running the ArchR function addTrajectory. Trajectory heatmaps for the peakMatrix, motifMatrix, and GeneIntegrationMatrix were then plotted using getTrajectory and plotTrajectoryHeatmap.

#### Survival analysis

Survival analysis on TCGA RNA-seq data was performed using the log-rank test and cox proportional hazard ratio in Gepia (Tang et al., 2017). Overall survival was used as the outcome to perform the statistical tests and patients were divided into two groups based on if expression of RUNX1 was in the top half or bottom half of patients.

#### Single-cell ATAC-seq: Construction of Normal Epithelial Reference and Projection into Normal Reference

To generate a normal epithelial reference, epithelial cells (as defined by gene activity scores of known epithelial marker genes) from 9 samples taken from 2 genetically normal donors were selected. Next, we followed a dimensionality reduction and clustering protocol similar to what is described above for the immune and stromal cell types. An iterative LSI dimensionality reduction was performed with addIterativeLSI with 4 iterations, 15,000 variable features, and clustering resolutions of 0.1, 0.1, and 0.2 following the first three iterations. Clusters were then defined with a resolution of 2 and a likely doublet cluster was removed, defined based on no clear accessibility around marker genes and the cluster consisting of cells from only a subset of the samples. The dimensionality reduction was then repeated with the above parameters. To simplify the clustering, we ran harmony batch correction (Korsunsky et al., 2019) on the IterativeLSI dimensionality reduction and clustered the cells with a resolution of 2.5. This did not substantially change the structure of the data, but facilitated cell type annotation. We note that cluster annotations would be similar without this step and we show all annotations on the UMAP generated from the IterativeLSI dimensionality reduction. Clusters were annotated based on gene activity scores at known marker genes. Marker genes include *DCLK1*, *HTR3C*, *HTR3E*, *B4GALNT4* for tuft cells; *KLK1*, *ITLN1*, *WFDC2*, and *CLCA1* for immature goblet cells; *MUC2*, *TFF1*, *FCGBP*, and *TBX10* for goblet cells; *CA1* for immature enterocytes, *RAB6B* for enterocytes; *CRYBA2* and *SCGN* for enteroendocrine cells; *BEST4*, *CA7*, *OTOP2*, and *OTOP3* for Best4+ enterocytes; and *SMOC2*, *RGMB*, *LGR5*, and *ASCL2* for stem cells. These marker genes are supported by the literature and many have been previously shown to be specific markers for these cell types in scRNA seq data (Smillie et al., 2019). ChromVar deviations for the normal epithelial reference were computed in ArchR as described above.

After generation of a normal epithelial reference, we next aimed to project diseased cells into this subspace, as has previously been done for placing diseased cells isolated from MPAL samples into the hematopoietic hierarchy (Granja et al., 2019). To accomplish this, we started with the tile matrix for epithelial cells from a given sample. We then selected the features from this tile matrix that were used in the final LSI iteration for the normal epithelial cells. We computed the inverse document frequency using the number of rows and columns from the initial LSI computation and performed the same svd transformation as was done in the final LSI iteration. After projecting each cell into this iterativeLSI subspace, we identified the 25 nearest neighbor cells with get.knnx in R. Cell type annotations were then assigned for each cell based on the most common cell type annotation of the 25 nearest neighbors.

#### Single-cell ATAC-seq: Peak calling for epithelial compartment

To generate a union peak set representing all samples and cell types present in the epithelial compartment, we wanted to ensure that we captured sample- and cell-type-specific peaks. To accomplish this, we divided the cells into groups, generated pseudobulk replicates for each group, called peaks on the pseudobulk replicates, generated a reproducible peak set for each group using the peaks called for the pseduobulk replicates, and then iteratively merged the peaks sets for each group into a union peak set using the approach previously described for scATAC data and implemented in ArchR (Granja et al., 2019, Granja et al., 2020). When selecting groups of cells for peak calling we choose not to create distinct groups for each epithelial cell type from each scATAC experiment because (A) this would result in a total of 979 groups (89 * 11 cell types) and (B) some of these groups would have very few cells and thus insufficient coverage to call peaks. To circumvent this, we opted for a mix of sample specific and cell type specific groupings. The following cell types consisted of a larger number of cells, so were initially divided into groups of each cell type from each sample: Stem, TA2, TA1, Enterocyte Progenitors, Immature Enterocytes, Enterocytes, Secretory TA, Immature Goblet, Goblet. The remaining cell types were divided into groups based on whether they originated from Normal, Unaffected, Polyp, or CRC samples (as defined by gross phenotype). After this initial classification, groups with fewer than 300 cells were identified. To preserve sample specific information, we first merged groups with their nearest cell type in the normal differentiation trajectory (e.g., if there were <300 enterocytes from a sample, that group was combined with the immature enterocyte group from the same sample). This was done iteratively with the following regrouping rules Enterocytes > Immature Enterocytes, Enterocyte Progenitors > Immature Enterocytes, TA1 > TA2, TA2 > Stem, Stem > TA2, Secretory TA > Immature Goblet, Goblet > Immature Goblet for groups with less than 300 cells prior to this step. For the cell type disease state groupings, there were insufficient cells for some of the enteroendocrine and Best4+ enterocyte groups, so normal and unaffected enteroendocrine groups were combined, polyp and CRC enteroendocrine groups were combined, and polyp and CRC Best4+ enterocyte groups were combined. After this step, we again identified any groups that did not have at least 300 remaining cells, and then combined the groups as follows:

1. Patient F Immature Enterocytes groups with fewer than 300 cells were merged into a single group.
2. Patient A002 Immature Enterocytes groups with fewer than 300 cells were merged into a single group.
3. Patient A014 Immature Enterocytes groups with fewer than 300 cells were merged into a single group.
4. Immature Goblet groups from A002 Unaffected samples with fewer than 300 cells were merged into a single group.
5. CRC-1-8810-Immature Goblet, CRC-2-15564-Immature Goblet, CRC-3-11773-Immature Goblet, and A001-C-007-Immature Goblet were merged together.
6. F091-Immature Goblet and F034-Immature Goblet groups were merged together.
7. F034-TA2 and F091-TA2 groups were merged together.
8. A015-C-001-Immature Goblet and A015-C-002-Immature Goblet groups were merged together.
9. A015-C-001-TA2 and A015-C-002-TA2 groups were merged together.
10. A014-C-008-Immature Goblet and A014-C-108-Immature Goblet groups were merged together.
11. A002-C-021-Immature Goblet and A002-C-016-Immature Goblet groups were merged together.

This process resulted in a total of 271 cell groupings with an average 1,327 cells per grouping. After defining these groupings, the ArchR functions addGroupCoverages followed by addReproduciblePeakSet were run using these groupings to group the cells for peak calling and setting all other parameters to their default values.

#### Single-cell ATAC-seq: Identification of Differential Peaks

To compute pairwise differential tests, the ArchR function getMarkerFeatures was used with testMethod set to wilcoxon and bias set to c(“TSSEnrichment”, “log10(nFrags)”) and useGroups and bgdGroups set to be the two samples being tested.

#### Single-cell ATAC-seq: Definition of Malignancy Continuum

To compute the malignancy continuum, we computed differential peaks between stem cells from all polyp, unaffected, and CRC samples in our dataset and normal stem cells. Unaffected samples from the same region of the colon in the same patient were merged together to ensure that there was a sufficient number of stem cells to compute differentials. After computing differentials individually for each sample, we selected the set of peaks that was significantly differential in at least two samples (Wilcoxon FDR≤0.05 and |Log_2_FC|≥1.5 in ≥2 samples). We then constructed the matrix of log_2_-fold changes for this set of significant peaks in all samples. The principal components of these differentials were computed with prcomp in R and a spline was fit to the first two principal components. For each sample, we then identified the nearest point on the spine (minimum euclidean distance), and the samples were ordered based on the position of the nearest point on the spline fit. We only included scATAC samples with at least 250 stem cells in the malignancy continuum, as we have higher confidence in the differentials computed with a larger number of cells.

#### Single-cell ATAC-seq: Identification of Differential Peaks and Enriched Motifs Along the Malignancy Continuum

To identify differential peaks along the malignancy continuum, unaffected samples were used as a background because we observed relatively small differences between unaffected and normal tissues, and because including more samples allowed us to better match potentially biasing features such as read depth and TSS enrichment when computing differentials. A few unaffected samples were found to have dysplasia on microscopic pathology and as a result were not included in the background when computing these differentials. We first identified differential peaks between stem-like cells in each polyp and CRC sample and stem cells from all unaffected tissues. Peaks that were significantly differential in at least two samples (Wilcoxon FDR≤0.05 and |Log_2_FC|≥1.5 in ≥2 samples) were clustered into 10 groups with kmeans clustering using the kmeans function in R with iter.max set to 500. Following clustering of the differential peaks into 10 groups, hypergeometric enrichment of motifs within those 10 groups was calculated using the ArchR function peakAnnoEnrichment. We note that this same analysis was done using normal stem cells as the background for differential testing and produced very similar results (Figures S4B and S4C).

#### Single-cell RNA sequencing

Initial processing of snRNA-seq data was done with the Cell Ranger Pipeline (https://support.10xgenomics.com/single-cell-gene-expression/software/pipelines/latest/what-is-cell-ranger) by first running cellranger mkfastq to demultiplex the bcl files and then running cellranger count. Since nuclear RNA was sequenced, data were aligned to a pre-mRNA reference.

#### Single-cell RNA-seq: Doublet removal and initial clustering

After running Cell Ranger, the filtered_feature_bc_matrix produced by Cell Ranger was read into R with the Seurat (Stuart et al., 2019) function Read10X. The data was filtered to remove cells with fewer than 400 unique genes per cell or greater than 4000 genes per cell. DoubletFinder (McGinnis et al., 2019) was run for each sample using PCs 1-20. nExp was set to 0.08*nCells^2^/10000, pN to 0.25, and pK to 0.09. Cells classified as doublets were then removed prior to additional analysis.

After running DoubletFinder, the remaining cells from all samples were merged into a single Seurat object and cells with greater than 10,000 counts/cell or greater than 5% mitochondrial RNA were removed. The data was then processed with Seurat’s standard pipeline. First, NormalizeData was run using the method LogNormalize and scale.factor of 10,000. Variable features were identified with Seurat’s findVariableFeatures using the vst method and 20,000 features. ScaleData was then run on all genes and PCs were computed with RunPCA. The cells were then clustered using Seurat’s FindNeighbors with dimensions 1-20 and FindClusters with a resolution of 1.0. Expression of marker genes in the resulting clusters was then used to label clusters as epithelial, stromal, or immune for downstream analysis.

#### Single-cell RNA-seq: Dimensionality Reduction, Clustering, and Annotation of Immune and Stromal Compartments

Cells from the immune and stromal subcompartments were analyzed with Seurat’s standard analysis pipeline. First, data was normalized with NormalizeData and scaled with ScaleData. Principal components were then computed on the scaled data and a UMAP was generated from the resulting PCs. Clusters were identified with the surat function FindNeigbors and FindClusters with a resolution of 0.5 for the immune cells and 0.5 for the stromal cells. Markers for each cluster were identified with the Seurat function FindMarkers. Low quality or likely doublet clusters were identified based on no expression of any marker genes or expression of marker genes from multiple cells types. After removal of low quality/doublet cluster, the dimensionality reduction and clustering was repeated on the resulting cells as described above but with a final clustering resolution of 2.1 for the immune cells and 1.0 for the stromal cells. Following clustering, immune and stromal clusters were annotated based on expression of known marker genes in each cluster. To support the immune annotations, the scRNA immune dataset was also annotated using SingleR (Aran et al., 2019). The HumanPrimaryCellAtlasData was used as a reference and was subsetted to include only reference data with the following main labels: “DC”, “Epithelial_cells”, “B_cell”, “Neutrophils”, “T_cells”, “Monocyte”, “Endothelial_cells”, “Neurons”, “Macrophage”, “NK_cell”, “BM”, “Platelets”, “Fibroblasts”, “Astrocyte”, “Myelocyte”, “Pre-B_cell_CD34-”, “Pro-B_cell_CD34+”, and “Pro-Myelocyte.” The cell type of each scRNA immune cell was then predicted with the SingleR function and labels set to label.fine. Following annotation, markers for each cell type were identified with the Seurat function FindAllMarkers.

#### Single-cell RNA-seq: Data Visualization

All Dot Plots were generated using the Seurat function DotPlot and all plots of expression on the UMAP projection were generated with the Seurat function FeaturePlot.

#### Single-cell RNA-seq: Construction of Normal Epithelial Reference and Projection into Normal Reference

To analyze the epithelial compartment, we first constructed a normal epithelial reference using epithelial cells from normal colon taken from genetically normal donors. We started with data normalized with Seurat’s normalizeData function. Next, the iterative LSI dimensionality reduction was computed using 4 total iterations, following a procedure outlined previously (Aran et al., 2019; Granja et al., 2019). For each iteration, the mitochondrial, ribosomal, and HLA genes were filtered out and, from the remaining genes, the top 1600 most variable genes were identified. We then computed the TF-IDF transformation on these genes and performed SVD on the transformed matrix, and provided dimensions 1–8 of this reduction as input to Seurat’s SNN clustering with resolution of 0.1. We summed the individual clusters single cells, computed the logCPM transformation with ‘edgeR::cpm(mat,log = TRUE,prior.count = 3),’ and then found the top 1600 variable genes across the clusters. A TF-IDF transformation was then computed on these variable genes and an SVD was then performed on the transformed matrix. Dimensions 1-8 were retained and clusters were identified using the Seurat functions findNeighbors and FInd Clusters, but with an increased resolution of 0.2. This process was repeated a total of 4 times with a resolution of 0.4 after the third iteration. After the final dimensionality reduction, we found that the 5th LSI component was highly correlated with the sample of origin (by biserial correlation), so removed that dimension and used dimensions 1-4 and 6-8 for additional downstream analysis.

This final LSI dimensionality reduction was provided as input to compute a UMAP representation of the data and the cells were clustered using a resolution of 1.0. The resulting clusters were then annotated based on expression of known marker genes. The projection of cells into the LSI subspace defined for normal colon epithelial cells was done following the procedure described previously (Granja et al., 2019). Briefly, when computing the TF-IDF transformation on normal colon epithelial cells, we stored the colSums, rowSums and SVD. To project cells from additional samples into this subspace, we first zero out rows based on the initial TF-IDF rowSums. We next calculated the term frequency by dividing by the column sums and computed the inverse document frequency from the previous TF-IDF transformation. These were then used to compute the new TF-IDF. The resulting TF-IDF matrix was projected into the previously defined SVD. Cells were classified by identifying their 25 nearest neighbors in the LSI subspace using get.knnx in R and then classifying the cell as the most common annotation for those 25 nearest neighbors.

#### Single-cell RNA-seq: Definition of Malignancy Continuum and Determination of Differential Genes Along Continuum

Similar to scATAC, differential genes were computed between stem cells from each sample and stem-cells from normal colon. To compute differential genes for the scRNA dataset, the Seurat function FindMarkers was used with ident.1 set as the sample of interest, ident.2 set as the background_sample, min.pct = 0, logfc.threshold = 0, min.cells.feature = 0, and max.cells.per.ident = 300.

For computing the snRNA-seq malignancy continuum, an analogous process to the one used for ATAC was carried out with the following minor differences: 1) Differential genes rather than differential peaks were used, 2) significance cutoffs for including a gene were wilcoxon p_adj_≤0.05 and |Log_2_FC|≥0.5 in ≥2 samples, and 3) we required there to be at least 100 snRNA cells in a group to compute differentials. Following determination of the RNA malignancy continuum, we computed differential genes between each polyp and CRC sample against all unaffected samples as was done for scATAC and plotted the set of differential genes with wilcoxon p_adj_≤0.05 and |Log_2_FC|≥0.75 in ≥2 samples in the heatmap in Figure 4.

### Analysis of DNA methylation data

Analysis of TCGA 450K methylation data was done with TCGABioloinks (Colaprico et al., 2016). Data for normal colon and colorectal adenocarcinoma were downloaded using the function GDCquery with project = c(”TCGA-COAD”), data.category = “DNA Methylation”, legacy = FALSE, platform = c(”Illumina Human Methylation 450”), and sample.type = c(”Primary solid Tumor”,”Solid Tissue Normal”). Differentially methylated probes between normal and colorectal adenocarcinoma samples were computed using TCGAanalyze_DMR with a p-value cutoff of 10^-5^ and a mean difference in β value cutoff of 0.25 to determine significance. Overlaps between DNA methylation probes and our peak set were identified with the GenomicRanges function FindOverlaps in R.

